# MiR-124 synergism with ELAVL3 enhances target gene expression to promote neuronal maturity

**DOI:** 10.1101/2020.03.25.006635

**Authors:** Ya-Lin Lu, Yangjian Liu, Matthew J. McCoy, Andrew S. Yoo

## Abstract

Neuron-enriched microRNAs (miRNAs), miR-9/9* and miR-124 (miR-9/9*-124), direct cell fate switching of human fibroblasts to neurons when ectopically expressed by repressing anti-neurogenic genes. How these miRNAs function after the onset of the transcriptome switch to a neuronal fate remains unclear. Here, we identified direct targets of miRNAs by Argonaute (AGO) HITS-CLIP as reprogramming cells activate the neuronal program and reveal the role of miR-124 that directly promotes the expression of its target genes associated with neuronal development and function. The mode of miR-124 as a positive regulator is determined by a neuron-enriched RNA-binding protein, ELAVL3, that interacts with AGO and binds target transcripts, whereas the non-neuronal ELAVL1 counterpart fails to elevate the miRNA-target gene expression. Although existing literature indicate that miRNA-ELAVL1 interaction can result in either target gene upregulation or downregulation in a context-dependent manner, we specifically identified neuronal ELAVL3 as the driver for miRNA target gene upregulation in neurons. In primary human neurons, repressing miR-124 and ELAVL3 led to the downregulation of genes involved in neuronal function and process outgrowth, and cellular phenotypes of reduced inward currents and neurite outgrowth. Results from our study support the role of miR-124 promoting neuronal function through positive regulation of its target genes.

## Introduction

MiR-9/9* and miR-124 (miR-9/9*-124) function as reprogramming effectors that, when ectopically expressed in human adult fibroblasts (HAFs), induce an extensive reconfiguration of the chromatin accessibility landscape leading to the erasure of fibroblast fate and activation of the neuronal program (Abernathy et al. 2017). The conversion process by miR-9/9*-124 shares similarities to molecular cascades underlying neurogenesis during neural development such as the downregulation of REST, a well-established transcription repressor of neuronal genes (Ballas et al., 2005; Lee et al., 2018; Schoenherr and Anderson, 1995) and switching of homologous chromatin modifiers from non-neuronal to neuronal counterparts including DNMT3B to DNMT3A, subunits of BAF/BRM-associated factor (BAF) complexes, and TOP2A to TOP2B (Abernathy et al., 2017a; Lee et al., 2018; Lessard et al., 2007; Staahl et al., 2013; Tsutsui et al., 2001; Watanabe et al., 1994; Yoo et al., 2009, 2011). Although the direct repression by brain-enriched miRNAs on non-neuronal targets for initiating the neuronal program (Makeyev et al., 2007; Packer et al., 2008; Visvanathan et al., 2007; Yoo et al., 2009) offer a classic representation of miRNAs acting as negative regulators of target genes, how these miRNAs function within the neuronal gene network remains unknown.

Previous studies demonstrated that miR-9/9*-124 expression is required throughout the conversion process till the endogenous miR-9/9* and miR-124 are activated (Abernathy et al., 2017a; Victor et al., 2014). Because early miRNA expression during neuronal conversion functions to repress non-neuronal targets expressed in fibroblasts, we reason that the miRNAs remain necessary for the induction of the neuronal program after the repression of anti-neurogenic genes. In the present study, we performed high-throughput sequencing of RNA isolated by crosslinking immunoprecipitation (HITS-CLIP) of Argonaute (AGO HITS-CLIP) during miR-9/9*-124-mediated neuronal reprogramming of human fibroblasts to map miR-9/9*-124-target interactions at the onset of neuronal fate acquisition which led to an unexpected identification of neuronal genes enriched with AGO binding, in particular, corresponding to the binding sites of miR-124. The upregulation of these neuronal genes requires miR-124, suggesting that miR-124 not only function as a repressor, but also as an effector to promote neuronal gene expression. Although the ability of miRNAs as a positive effector of downstream target genes has been implicated before (Truesdell et al., 2012; Vasudevan et al., 2007; Vasudevan and Steitz, 2007), little is known about the molecular mechanism that governs a miRNA’s activity as a positive regulator of target genes in neurons and the identity of activated target genes. Here, we use homologous genes, *PTBP1* and *PTBP2*, as a model to dissect how miR-124 selectively upregulates PTBP2 expression in neurons and reveal genetic networks that are enhanced by miR-124 in both reprogrammed and primary human neurons.

## Results

### AGO binds both downregulated and upregulated genes during miR-9/9*-124-mediated direct neuronal reprogramming of human fibroblasts

Previous studies showed that the ectopic expression of miR-9/9*-124 in human adult fibroblasts (HAFs) induces a neuronal state characterized by the appearance of neuronal markers (such as MAP2, TUBB3, NCAM, and SNAP25) (Figure 1A) and electrical excitability (Abernathy et al. 2017). To identify target genes of miR-9/9*-124 as reprogramming cells transition to neuronal identity, we carried out AGO HITS-CLIP after two weeks into reprogramming, a time point when neuronal genes are activated (Abernathy et al. 2017), to identify transcripts bound with AGO loaded with miR-9/9* or miR-124 over the non-specific miRNA (miR-NS) control Log_2_FC ≥ 1; adj.P-value < 0.05). We compared these hits to the list of differentially expressed genes (DEGs) (by RNA-seq analysis; −1 ≤ Log_2_FC ≥ 1; adj.P-value < 0.05) at day 20 of miRNA-induced neurons (miNs) (Abernathy et al. 2017). As expected, we found target transcripts that were downregulated consistent with the repressive mode of miRNAs (Figure 1B, 113 genes labeled as green dots) including some known targets, *SHROOM3* and *PHF19* (Lu et al., 2018; Neo et al., 2014; Zhou et al., 2014). Interestingly, we also discovered 453 unique gene transcripts with enriched AGO binding, which were upregulated in day 20 miNs (Figure 1B, red dots). By gene ontology (GO) analysis, these upregulated genes with enriched AGO loading in response to miR-9/9*-124 were associated with various neuronal processes such as synaptic transmission and regulation of membrane potential (red), in contrast to non-neuronal, downregulated target genes (green) (Figure 1C). Moreover, when examined against the time-course transcriptome analysis during neuronal reprogramming (Abernathy et al. 2017), the expression of the identified neuronal target genes showed continuous upregulation during later time point of neuronal conversion (Figure 1D). For example, neuronal transcripts such as *MAP2, PTBP2*, and *SLC4A8* that were highly expressed in day 20 miNs harbored AGO binding sites at the 3’UTR (Figure 1E). To determine the fraction of upregulated neuronal genes that are likely actual targets of miR-9/9* and/or miR-124, we extracted the AGO-enriched sequences and predicted the duplex formation by either miR-9/9* and/or miR-124 through RNAhybrid (a maximum free energy threshold of −20 kcal/mol) (Rehmsmeier, 2004). Of the 453 upregulated DEGs in miNs bound by AGO (Figure 1B), 328 (∼72%) gene transcripts are predicted to contain miR-9/9* and/or miR-124 binding sites (Figure 1F). These transcripts harboring miR-9/9*-124 sites are also associated with the similar set of neuronal GO terms in Figure 1C as they include a selection of genes important for neuronal function (Figures 1F). Of these upregulated miR-9/9*-124 target genes, more than 94% of the total genes (308 of 328 genes) contain miR-124 target sequence alone (45%) or with miR-9/9* sites, while less than 6% of the genes (20 genes) contain miR-9/9* target sites only (Figure 1G). These results collectively demonstrate that the AGO-loaded transcripts differentially (up or down) respond to miR-9/9*-124 during neuronal conversion.

**Figure 1.**
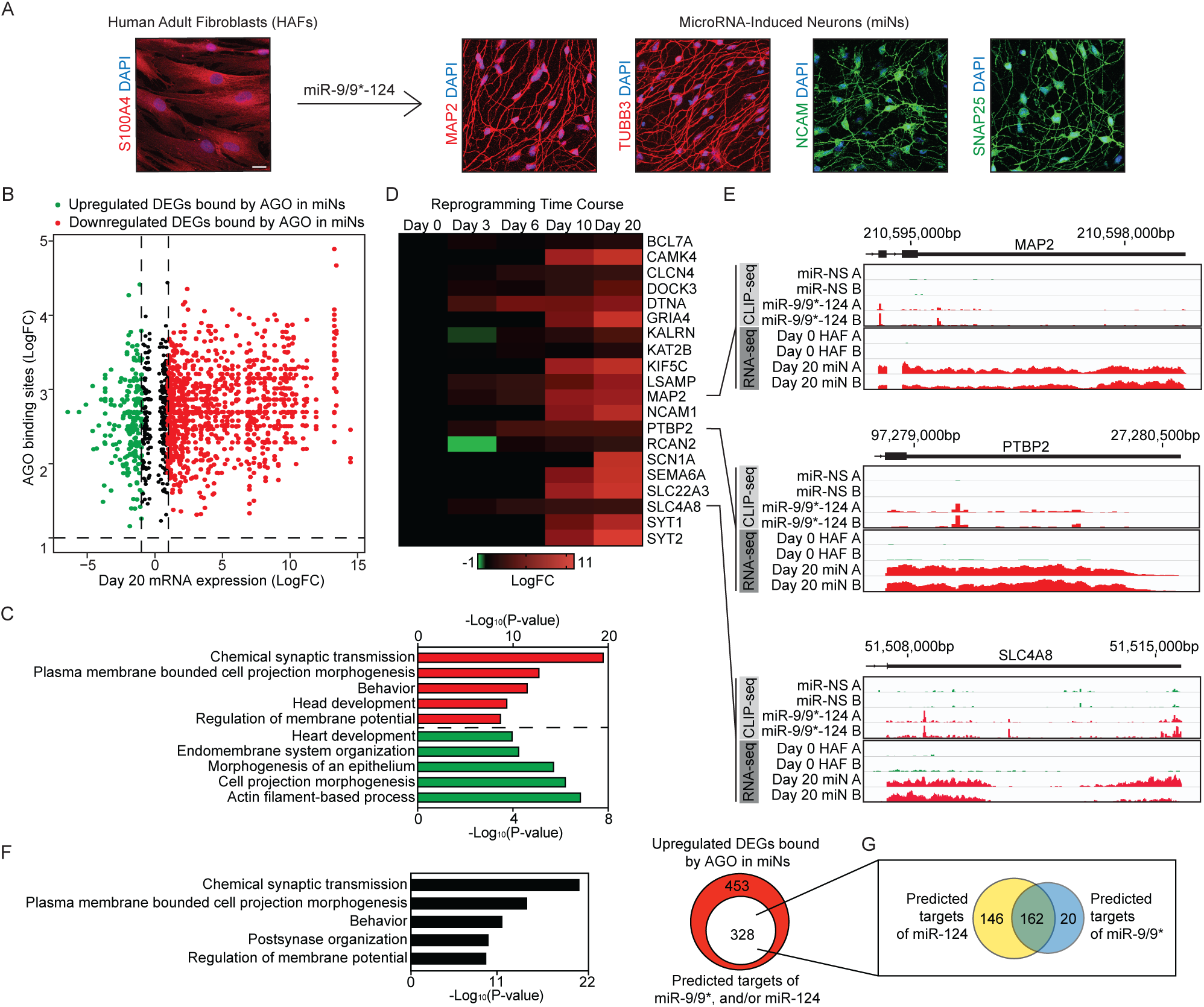
Enrichment of AGO binding on transcripts of neuronal genes upregulated during miR-9/9*-124-mediated neuronal conversion. (A)Examples of miR-9/9*-124-mediated direct reprogramming of human adult dermal fibroblasts (HAFs) into neurons (miNs). Photographs show immunostaining of the fibroblast marker (S100A4) and neuronal markers (MAP2, TUBB3, NCAM, and SNAP25). Scale bar = 20 μm. (b)A volcano plot of differentially expressed genes at day 20 miNs and enrichment of AGO binding in response to miR-9/9*-124 expression identified by AGO HITS-CLIP analysis. Red dots, day 20 mRNA expression Log2FC ≥ 1; adj.P-value < 0.05, AGO binding Log2FC ≥ 1, adj.P-value < 0.05. Green dots, day 20 mRNA expression Log2FC ≤ −1; adj.P-value < 0.05, AGO binding Log2FC ≥ 1, adj.P-value < 0.05. (c)Top biological GO terms of upregulated (red) and downregulated (green) DEGs in day 20 miNs differentially bound by AGO. (D)Time course heatmap of select upregulated DEGs enriched for AGO binding with miR-9/9*-124 expression. (E)Track views of AGO CLIP-seq tracks (top) and RNA-seq tracks (bottom) for gene examples showing the enrichment of AGO binding miR-9/9*-124 expression (over control miR-NS) and increased transcript levels at 3’UTRs. (F)Of the 301 unique upregulated DEGs identified in (A), 207 are predicted to harbor miR-9/9* and/or miR-124 sites at the AGO-enriched regions through RNAhybrid prediction. The graph indicates top biological GO terms associated with the 328 upregulated DEGs containing miR-9/9* and/or miR-124 target sites. (G)Breakdown of the 328 upregulated DEGs in (F) based on common or specific targets of miR-9/9* and/or miR-124.

### MiR-124 target genes are upregulated during neuronal conversion

As a large fraction of the upregulated genes contained miR-124 target sequences (Figure 1G), we further tested if the upregulated genes were *bona fide* targets of miR-124 by knocking down

miR-124 expression through the use of a tough decoy (TuD) to inhibit miRNA activity (Bak et al., 2013; Haraguchi et al., 2009). The effect of the lentivirus-based TuD for miR-124 (TuD-miR-124) was monitored after transduction by following the concurrent expression of TurboRFP reporter built in the lentiviral vector and measuring the mature miR-124 level. TuD-miR-124 yielded more than 60% reduction of miR-124 expression in comparison to the control, non-specific miRNA tough decoy (TuD-miR-NS) (Figure S1A-S1B). We then performed RNA-seq analysis on day 20 miNs treated with either TuD-miR-NS or TuD-miR-124 to see whether any of the miR-124 target would fail to be upregulated with TuD-miR-124 treatment (Figure 2A, S1C). When compared with upregulated DEGs in day 20 miNs that were also enriched for AGO binding (Figure 1), we identified 192 genes (Log_2_FC ≤ 0.5; adj.P-value < 0.01) that failed to be upregulated upon miR-124 reduction (Figure 2A, Table S1). With GO analysis, these identified genes were associated with neuronal terms involved in various synaptic processes (Figure 2B).

**Figure 2.**
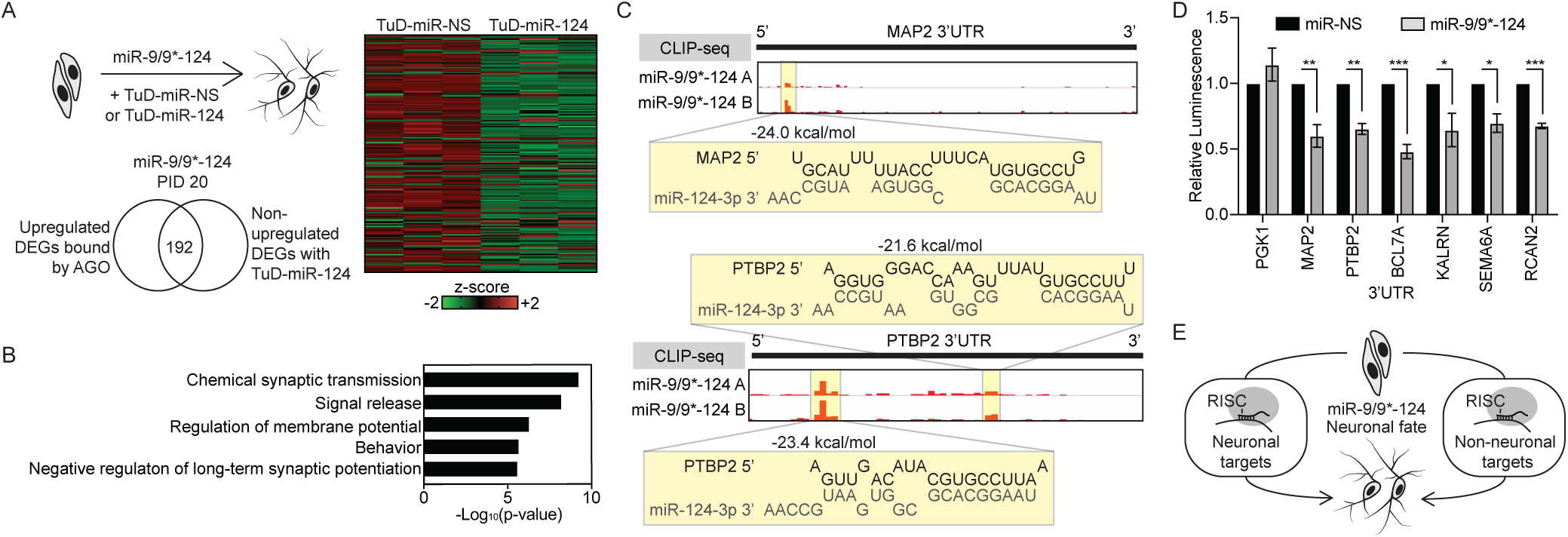
Identification of miR-124 target genes that fail to be upregulated upon the inhibition of miR-124 during miRNA-mediated neuronal conversion. (A)Left, miR-124 activity is reduced through the use of tough decoy (TuD). By overlapping with upregulated DEGs bound by AGO (Figure 1), 192 genes were identified as genes that fail to be increased upon the reduction of miR-124. Right, a heatmap of z-scores of the 192 genes from RNA-seq comparing miNs between TuD-miR-NS and TuD-miR-124 treatments. (B)Top biological GO terms associated with the 192 genes identified in (A). (C)Predicted binding of miR-124 to the 3’UTR sequences of neuronal target genes, for example, *MAP2* and *PTBP2*, according to RNAhybrid prediction at the highlighted AGO HITS-CLIP peaks. (D) Luciferase assays in HEK293T cells of upregulated neuronal target genes of miR-124 selected from A). In the non-neuronal context of HEK293T cells, 3’UTRs from the neuronal genes are targeted and repressed, instead, by miR-9/9*-124. Luminescence measured after 48 hrs of transfection and normalized to miR-NS control of each condition. Data are represented by mean ± SEM from three independent experiments (from left, ** P = 0.0094, P = 0.0011; *** P < 0.001; * P = 0.048, P = 0.012; *** P < 0.001). (E)A diagram of the observed phenomenon in which miR-9/9*-124 can promote neuronal identity by simultaneously targeting both non-neuronal genes for repression while promoting the expression of neuronal genes during neuronal conversion.

To further analyze miR-124-target interactions, we looked into the potential duplex formations between miR-124 and the identified AGO HITS-CLIP peak regions using RNAhybrid (a maximum free energy threshold of −20 kcal/mol) (Rehmsmeier, 2004). Of the upregulated transcripts that harbor AGO-enriched peaks, we identified a spectrum of miR-124 and target mRNA duplex configurations ranging from the canonical 2-8 seed base pairing to non-canonical base pairing starting at position 3 and 4 (Figures 2C, S1D, and S2). These examples include *MAP2* 3’UTR, in which miR-124 base pairing is predicted to start at position 3, while miR-124 target sequences on *PTBP2* 3’UTR at the two peaks are predicted to start at position 1 and position 2, respectively (Figure 2C). Interestingly, across the predicted miR-124:target duplex configurations, we observed consistent auxiliary 2-3 base pairing at the 3’ end of miR-124 (Figure S1D).

Our results so far indicate that miR-124 can target and promote the expression of select neuronal genes when cells acquire the neuronal fate during neuronal conversion. We wondered if the active mode of miR-124 would be specific to neuronal cell types. We cloned the 3’UTRs of several identified, upregulated targets (*MAP2, PTBP2, BCL7A, KALRN, SEMA6A, RCAN2*) into a luciferase reporter construct and transfected into a non-neuronal cell type, HEK293T (a human embryonic kidney cell line), with a miR-9/9*-124 expression construct. After 48 hours post-transfection, we found that unlike reprogrammed neurons, miR-9/9*-124 instead, repressed the luciferase signal in HEK293T cells in comparison to the control miR-NS, while the non-target control, *PGK1* 3’UTR was unaffected (Figure 2D). This result suggests that additional determinants available in neuronal cells may be in play to govern the activity of miR-124 as a positive regulator while simultaneously functioning to repress non-neuronal targets (Figure 2E).

### PTBP1 and PTBP2 3’UTR as targets of miR-124

To further dissect the mechanism underlying the dual modes of miR-124 on its targets, we elected to focus on *PTBP2* from the list of our identified neuronal targets because i) *PTBP2* and its non-neuronal homolog, *PTBP1*, both contain miR-124 sites in their 3’UTRs (Figure 3A), and ii) both *PTBP1* and *PTBP2* 3’UTRs are targeted and repressed by miR-124 in HEK293T cells (Figure 3A), and iii) contrastingly, as observed in the human brain, qPCR analysis showed that the neuronal conversion established the mutually exclusive expression between *PTBP1* and *PTBP2* (Figure 3B). Therefore, the PTB homologs represent an ideal example to investigate how two closely related targets respond differently to miR-124.

**Figure 3.**
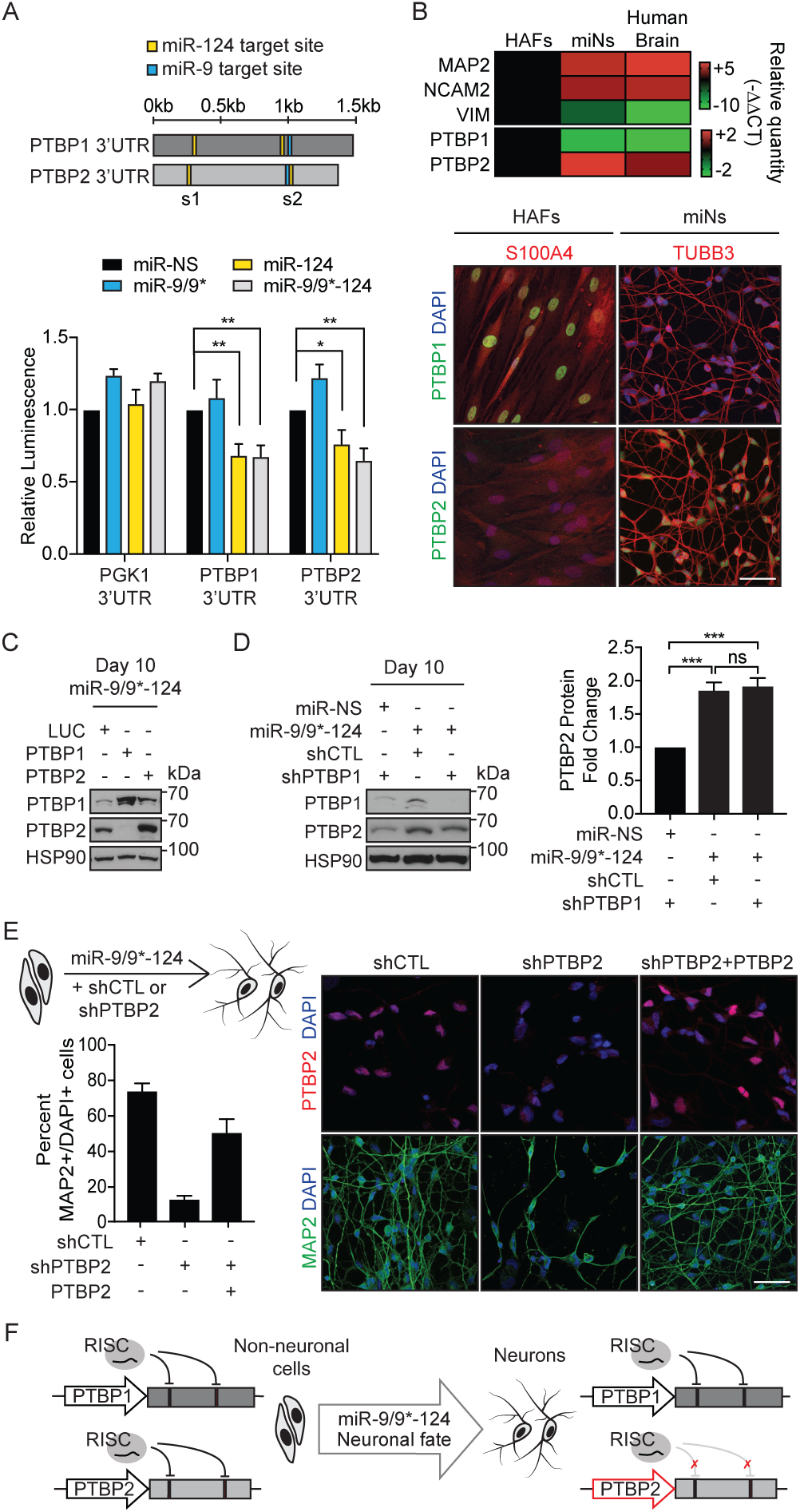
MiR-124 targets both PTBs, but differentially regulate PTB expression during neuronal conversion. (A)Top, a schematic diagram of PTB 3’UTRs with miR-124 (yellow) and miR-9 (blue) target sites. S1 and s2 refer to the two conserved miR-124 target sites on PTBP2 3’UTR. Bottom, luciferase assays with luminescence measured after 48 hrs of transfection and normalized to miR-NS control in each condition. Data are represented by mean ± SEM from four independent experiments. Two-way ANOVA followed by Dunnett’s test (from left, PTBP1 3’UTR ** P = 0.0043, 0.0034; PTBP2 3’UTR * P = 0.0355, ** P = 0.0016) (B)Top, A heatmap of gene expression assessed by qPCR in starting HAFs, day 30 miRNA-induced neurons (miNs), and human brain RNA. Bottom, PTB switching is recapitulated during the miRNA-mediated direct conversion of HAFs into neurons. HAFs and day 30 miNs immunostained for PTBP1 and PTBP2, along with a fibroblast marker, S100A4, and a pan-neuronal marker, TUBB3. Scale bar = 50 μm. (C)Initial induction of PTBP2 by the reduction of PTBP1 by either shRNA or miRNAs during miRNA-mediated neuronal reprogramming. PTBP1 overexpression in the miR-9/9*-124 expression background suppresses the PTBP2 induction. (D)Left, an immunoblot showing that PTBP1 knockdown in HAFs resulted in induction of PTBP2, but PTBP2 expression becomes more pronounced in the presence of miR-9/9*-124 compared to PTBP1 knowdown only. Right, quantification of PTBP2 band intensity as relative fold changes compared to PTBP1 knockdown alone. Data were normalized to HSP90 from four independent experiments. The plots were represented in mean ± SEM. One-way ANOVA followed by Tukey’s test (from top *** P = 0.0002, 0.0004; ns P = 0.8925). (E)Top left, a schematic diagram of the experimental procedure. Right, photographs of day 30 miNs treated with CTL shRNA, PTBP2 shRNA, or PTBP2 shRNA with PTBP2 cDNA. Cells were immunostained for PTBP2 and MAP2. Bottom left, quantification of the percentage of MAP2-positive cells with two or more neurite processes over the total number of DAPI-positive cells. Scale bar = 50 μm. Data are represented as mean ± SEM. One-way ANOVA followed by Tukey’s test (*** P < 0.0001; ** P 0.0016). shCTL n = MAP2 356/482; shPTBP2 n = MAP2 31/239; shPTBP2+PTBP2 n = MAP2 225/454. (F)A model of differential miR-124 activity on PTB 3’UTRs during neuronal conversion.

The 3’UTRs of human *PTBP1* and *PTBP2* contain two predicted target sites for miR-124 (yellow bar) and one site for miR-9 (blue bar) (Figure 3A). Luciferase reporter assays in the non-neuronal HEK293T cells showed that miR-9/9*-124 or miR-124 alone (but not miR-9/9* alone) repressed both *PTBP1* and *PTBP2* through their 3’UTRs indicating that miR-124 is the primary miRNA targeting both PTB 3’UTRs (Figure 3A). Mutating the two miR-124 sites in *PTBP2* 3’UTR (s1 and s2) rendered the 3’UTR insensitive to miR-124 (Figure S3A). While our results pointing to the ability of miR-124 to target and repress 3’UTRs of both *PTBP1* and *PTBP2* in cell lines were consistent with previous findings (Makeyev et al., 2007; Xue et al., 2016), it remains unknown how miRNA-mediated neuronal conversion of HAFs establishes the mutually exclusive expression of PTBP1 and PTBP2 as seen in the human brain (Figure 3B) and in *in vivo* neurons (Boutz et al., 2007).

Because *PTBP1* and *PTBP2* are regulated differentially in miNs, we asked whether the repressive activity of miR-124 on *PTBP2* 3’UTR could be reversed with a prolonged neurogenic input by miR-9/9*-124 in HEK293T cells. We expressed a destabilized EGFP reporter containing *PTBP1* 3’UTR, *PTBP2* 3’UTR, or control 3’UTR (CTL) that lacks a 3’UTR in HEK293T cells with the miR-9/9*-124 expression construct (Figure S3B). Analogous to PTBP2 upregulation during neuronal reprogramming of HAFs, we observed the selective repression of EGFP with *PTBP1* 3’UTR but not *PTBP2* 3’UTR (Figures S3B-C). These results suggest that with the prolonged neurogenic input, *PTBP2* 3’UTR responds to miR-124 differentially from when measured after 48 hours with miRNA expression in HEK293T cells (Figures 2A and 3A).

### MiR-124 accentuates PTBP2 expression beyond the induction mediated by the downregulation of PTBP1

Previous studies have shown that during development, PTBP1 destabilizes *PTBP2* transcript in non-neuronal cells by alternative splicing, and at the onset of neurogenesis, miR-124 directly represses *PTBP1* resulting in PTBP2 induction (Boutz et al., 2007; Makeyev et al., 2007). Upon miR-9/9*-124 expression in HAFs, we observed the concomitant downregulation and upregulation of endogenous PTBP1 and PTBP2, respectively (Figure 3B). As expected, the initial PTBP2 upregulation was abrogated when *PTBP1* cDNA was overexpressed (Figure 3C), supporting the role of PTBP1 reduction as the initiation step of PTBP2 expression. We then sought to stratify the contribution of PTBP1 repression alone versus the input of miRNAs to the overall PTBP2 level. When PTBP2 protein levels were compared between PTBP1 knockdown with a short hairpin RNA (shRNA) and miR-9/9*-124 expression, we found that miR-9/9*-124 enhanced the PTBP2 level by approximately a two-fold increase over the PTBP1 knockdown alone condition (Figure 3D). In fact, PTBP2 was more upregulated with miR-9/9*-124 despite the more pronounced reduction of PTBP1 with shRNA (Figure 3D), demonstrating that miR-9/9*-124 accentuate PTBP2 expression beyond the level induced by PTBP1 downregulation.

### Functional significance of PTBP2 upregulation for neuronal conversion

To probe the functional importance of *PTBP2* expression, we knocked down *PTBP2* with an shRNA during neuronal conversion. *PTBP2* shRNA completely impaired the induction of neurons marked by the loss of MAP2 expression, a neuronal marker (Figure 3E). The effect of shRNA was specific to *PTBP2* knockdown as supplementing *PTBP2* cDNA rescued the reprogramming defect (Figure 3E). These findings were somewhat surprising as the results from a previous study indicated that sequential reduction of PTBP2 by shRNA was found to promote maturation of reprogrammed neurons (Xue et al. 2016); however, our results indicate that at least at the initiation of reprogramming, PTBP2 is critical. While it is not clear why shRNA-versus miRNA-based reprogramming approaches lead to these different results, our results demonstrate the essential role of PTBP2, at least, at the onset of neuronal conversion. Furthermore, knocking down PTBP2 in primary cultured human neurons resulted in increased cell death as measured with SYTOX assay (Figure S4A-B), consistent with previous studies that showed the essential function of PTBP2 in primary neurons (Li et al., 2014).

We also performed Human Clariom D Assay in HAFs expressing the non-specific control miR-NS, miR-9/9*-124, or miR-9/9*-124 with *PTBP2* shRNA to identify genes whose expression and alternative splicing patterns were affected by the reduction of PTBP2. By two weeks, miR-9/9*-124-expressing cells showed significant downregulation of *PTBP1* and other fibroblast-enriched genes such as *FBN1* and *S100A4*, and upregulation of *PTBP2* and neuronal genes (for example, *NEFM*, and *SNAP25*) in comparison to the control miR-NS (FC ≥ 1.5, ANOVA p-value < 0.05) (Figure S4C) (Abernathy et al., 2017). Alternative splicing events mediated by PTBP2 were examined by comparing spliced events between *PTBP2* shRNA and control shRNA (shCTL) conditions. We found that reducing PTBP2 led to changes in splicing events (−2 ≤ splicing index ≥ 2, ANOVA p-value < 0.05) indicated by red (positive splicing index) and green dots (negative splicing index) (Figure S4D). This analysis identified known splicing targets of PTBP2, such as the exon skipping or exclusion in *UNC13B, DLG4*, and *CADM3*, and inclusion in *DNM1* and *SMARCC2* transcripts (Figure S4D) (Li et al., 2014; Licatalosi et al., 2012; Vuong et al., 2016; Zheng et al., 2012). Comparing genes upregulated in response to miR-9/9*-124 to genes associated with PTBP2-mediated alternative splicing (differential splicing events between miR-9/9*-124-shCTL and shPTBP2 conditions), identified 1183 differentially spliced transcripts of genes involved in processes such as neuronal differentiation and signaling (Figure S4E) (Table S2). Altogether, our results support the role of PTBP2 as a crucial regulator of the neuronal program.

### Differential sequence composition between PTBP1 and PTBP2 3’UTRs

Using PTBP1 downregulation and PTBP2 upregulation as a model, we sought to identify effectors that determine miR-124 function as a positive regulator. RNA-binding proteins (RBPs) have been shown to interact with miRNA-loaded RISC complexes to modulate target gene expression (Iadevaia and Gerber, 2015; Jiang and Coller, 2012; Plass et al., 2017). We ran the sequence of *PTBP2* 3’UTR through three RBP motif prediction databases, including RBPDB (Cook et al., 2011), RBPmap (Paz et al., 2014), and beRBP (Yu et al., 2018) (Figure S5A). Across the three databases, two RBPs, ELAVL1 and PUM2, were consistently predicted to bind to *PTBP2* 3’UTR (Figure S5A). As PTBP2 upregulation occurs in neurons, we focused on the family of RBPs whose expression is neuronally enriched. Whereas *ELAVL1* and Pumilio family RBPs, *PUM2* or its homolog, *PUM1*, are ubiquitously expressed (Lin et al., 2018; Okano and Darnell, 1997; Spassov and Jurecic, 2002), other ELAVL family members, *ELAVL2, ELAVL3*, and *ELAVL4* (collectively referred to as neuronal *ELAVLs, nELAVLs*), have been shown to be neuronally enriched (Okano and Darnell, 1997). We also examined the HITS-CLIP data of nELAVLs in the human brain (Scheckel et al., 2016) and found nELAVLs to be highly enriched at *PTBP2* 3’UTR, in contrast to *PTBP1* 3’UTR (Figure S5B-C). Interestingly, *PTBP2* 3’UTR contains AU-rich elements (AREs) that ELAVLs have been shown to bind to (Ince-Dunn et al., 2012; Scheckel et al., 2016), in contrast to *PTBP1* 3’UTR that lacks the ARE (Figure S5B). Moreover, nELAVL binding mapped to the ARE around the first miR-124 site (s1; seed, highlighted in red) *PTBP2* 3’UTR (Figure S5C). We thus examined by qPCR if nELAVLs are induced during neuronal conversion as well as other brain-enriched RBP markers including NOVA-, RBFOX-family proteins, and SSRM4. We found selective upregulation of nELAVLs with other neuronal RBPs in miNs similarly to the human brain, in contrast to the ubiquitous expression of *ELAVL1* (Figure 4A) (Okano and Darnell, 1997). To determine if nELAVL induction occurs concurrently with PTBP2 upregulation, we assessed ELAVL expression at multiple time points of neuronal conversion. We found that the transcriptional activation of nELAVLs, with *ELAVL3* being the most robust one (blue), aligned with the upregulation of *PTBP2* (black) by nine days into reprogramming (Figure 4B).

**Figure 4.**
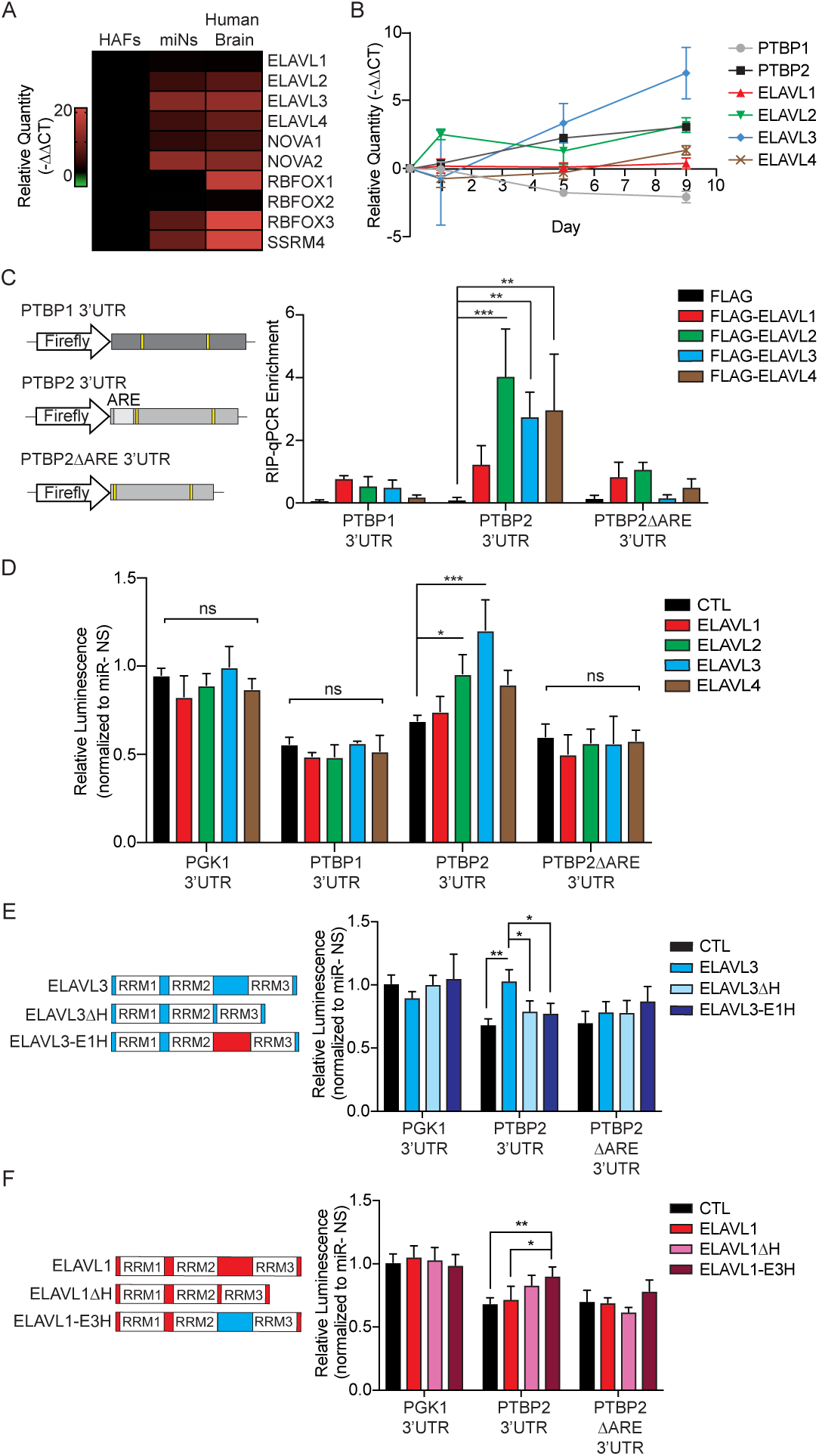
MiRNA-mediated PTBP2 upregulation requires nELAVL binding at PTBP2 3’UTR. (A)An expression heatmap of neuronal-enriched RBPs determined by qPCR in starting HAFs, day 20 miNs, and human brain RNA. (B)Time course qPCR assays of PTB and ELAVL transcripts during neuronal reprogramming. (C)Left, a diagram of luciferase constructs containing different PTB 3’UTRs: PTBP1 3’UTR, and PTBP2 3’UTR with and without (PTBP2ΔARE) the AU-rich element (ARE) used for RIP and luciferase assays. Right, RIP of individual FLAG-tagged ELAVLs 48 hrs after transfection. Enrichment normalized to input determined by qPCR for PTBP1 or PTBP2 3’UTR. Data are represented by mean ± SEM from three independent experiments. Two-way ANOVA followed by Dunnett’s test (from left, PTBP2 3’UTR *** P = 0.0001, ** P = 0.0086, 0.0042). (D)Luciferase assays with the addition of individual ELAVLs in the luciferase constructs containing the control PGK1, PTBP1, PTBP2 or PTBP2ΔARE 3’UTR. Luminescence was measured 48 hrs after the transfection and normalized miR-9/9*-124 to miR-NS control of each condition. Data are represented by mean ± SEM from at least three independent experiments. Two-way ANOVA followed by Dunnett’s test (* P = 0.0312, *** P = 0.0001). (E)Left, a schematic diagrm of ELAVL3 hinge mutants. Right, luciferase assays with the addition of wild-type ELAVL3 or ELAVL3 hinge mutants (ELAVL3Δ H: ELAVL3 hinge deletion; ELAVL3-E1H: ELAVL3 with ELAVL1 hinge). Luminescence was measured 48 hrs after transfection and normalized miR-9/9*-124 to miR-NS control of each condition. Data are represented by mean ± SEM from at least four independent experiments. Two-way ANOVA followed by Tukey’s test (from left ** P = 0.0020, * P = 0.0435, * P = 0.0286). (F)Left, a schematic diagram of ELAVL1 hinge mutants. Right, luciferase assays with the addition of wild-type ELAVL1 or ELAVL1 hinge mutants (ELAVL1 ΔH: ELAVL1 hinge deletion; ELAVL1-E3H: ELAVL1 with ELAVL3 hinge). Luminescence was measured 48 hrs after transfection and normalized miR-9/9*-124 to miR-NS control of each condition. Data represented in mean ± SEM from at least four independent experiments. Two-way ANOVA followed by Tukey’s test (** P = 0.0069, * P = 0.0260).

### MiRNA-mediated PTBP2 induction requires ELAVL3 binding at PTBP2 3’UTR

We then tested whether ARE within *PTBP2* 3’UTR would serve as a sequence that binds ELAVL proteins by expressing 3’UTRs of *PTBP1, PTBP2*, or *PTBP2* without ARE (*PTBP2ΔARE* 3’UTR) with individual FLAG-tagged ELAVLs (ELAVL1-4) in HEK293T cells (Figure 4C). RNA-immunoprecipitation (RIP) of FLAG-ELAVLs with FLAG antibody, followed by qPCR for detecting the loaded 3’UTRs using primers specific for either *PTBP1* or *PTBP2* displayed significant enrichment for *PTBP2* 3’UTR with nELAVL (ELAVL2, 3, and 4) pull-downs, while the binding of ELAVL1 to *PTBP2* 3’UTR was minimal. We could not detect significant enrichment for *PTBP1* 3’UTR for any of the ELAVLs (Figure 4C). Importantly, deleting ARE in *PTBP2* 3’UTR (*PTBP2ΔARE* 3’UTR) abolished the binding of nELAVL proteins to *PTBP2* 3’UTR (Figure 4C) indicating that ARE within *PTBP2* 3’UTR serves to recruit nELAVLs.

We then asked whether adding nELAVLs to the non-neuronal context of HEK293T cells, thereby reconstituting nELAVLs that become available during neuronal conversion, would alleviate miR-124-mediated repression of PTBP2 and enhance PTBP2 expression. Adding individual ELAVLs to control *PGK1* 3’UTR had no effect on luminescence (relative luminescence of ∼ 1; *PGK1* 3’UTR histograms) (Figure 4D). Luciferase activities with *PTBP1* 3’UTR remained repressed upon miR-9/9*-124 expression compared to the control miR-NS irrespective of the ELAVL addition (relative luminescence of < 1; *PTBP1* 3’UTR histograms) (Figure 4D). MiR-9/9*-124 led to the repression *PTBP2* 3’UTR in the absence of ELAVLs (black, relative luminescence ratio < 1; *PTBP2* 3’UTR histograms) (Figure 4D). However, adding ELAVL2 (green) and ELAVL3 (blue) significantly alleviated miR-9/9*-124-mediated repression on *PTBP2* 3’UTR, with especially ELAVL3 (blue) having the most significant effect on elevating the luciferase activity in comparison to the control construct (Figure 4D). To examine the requirement of ARE, we repeated the ELAVL addition experiments using the luciferase cassette with *PTBP2* 3’UTR lacking the ARE sequence (*PTBP2ΔARE*). Deleting ARE abolished the effect of ELAVL3 (blue) on *PTBP2* (*PTBP2ΔARE* 3’UTR histograms), which stayed repressed (Figure 4D), demonstrating the requirement of ELAVL3 and ARE for *PTBP2* upregulation by miR-124.

### Selective activity of ELAVL3 on PTBP2 3’UTR is dependent on the hinge region

ELAVL1-4 members exhibit high sequence homology across all three functional RNA recognition motifs (RRMs) except for the non-conserved spacer region (also referred to as hinge region) flanked by RRM2 and RRM3 (Hinman et al., 2013; Okano and Darnell, 1997). To better understand the specificity of ELAVL3 on *PTBP2* 3’UTR regulation, we mutagenized the hinge region of *ELAVL1* and *ELAVL3* by deleting or swapping the hinge region (H) between *ELAVL1* and *ELAVL3*. Deleting the H in *ELAVL3* (*ELAVL3ΔH*) (light blue) abrogated the alleviating effect on *PTBP2* 3’UTR repression, whereas no effect was observed with *PGK1* 3’UTR (Figure 4E, *PGK1* 3’UTR histograms). Moreover, replacing the *ELAVL3* H with *ELAVL1* H (*ELAVL3-E1H*) (dark blue) led to the failure of alleviating the *PTBP2* 3’UTR repression (Figure 4E), and none of the *ELAVL3* variants (wild-type and mutants) had any effect on *PTBP2ΔARE* 3’UTR (Figure 4E, *PTBP2ΔARE* 3’UTR histograms). These results indicate that the specificity of ELAVL3 to *PTBP2* 3’UTR is mediated by the *ELAVL3* H region. This notion is further supported by the increase in the luminescence readout of *PTBP2* 3’UTR in HEK293T cells when *ELAVL1* H is replaced by *ELAVL3* H (*ELAVL1-E3H*) (dark red) compared to wild-type *ELAVL1* (red) (Figure 4F, *PTBP2* 3’UTR histograms).

### ELAVL3 promotes PTBP2 expression during neuronal reprogramming

To further examine if ELAVL3 would be critical for PTBP2 upregulation during the neuronal conversion of HAFs, we knocked down *ELAVL3* by shRNA (shELAVL3) to assess PTBP2 expression and neuronal reprogramming. Knocking down ELAVL3 resulted in the significant downregulation of PTBP2 expression as determined by immunostaining, qPCR, and immunoblotting analyses, and impairment of the conversion process (Figures 5A-C). This knockdown effect was specific for ELAVL3 downregulation as PTBP2 expression and neuronal fate acquisition could be rescued by overexpressing *ELAVL3* cDNA in the presence of shELAV3 (Figure 5A). It is noteworthy that reducing the function of other nELAVLs (ELAVL2 and 4) had a milder effect on PTBP2 expression (Figures 5B-C), highlighting the role of ELAVL3 as a primary driver for PTBP2 upregulation with miR-124.

**Figure 5.**
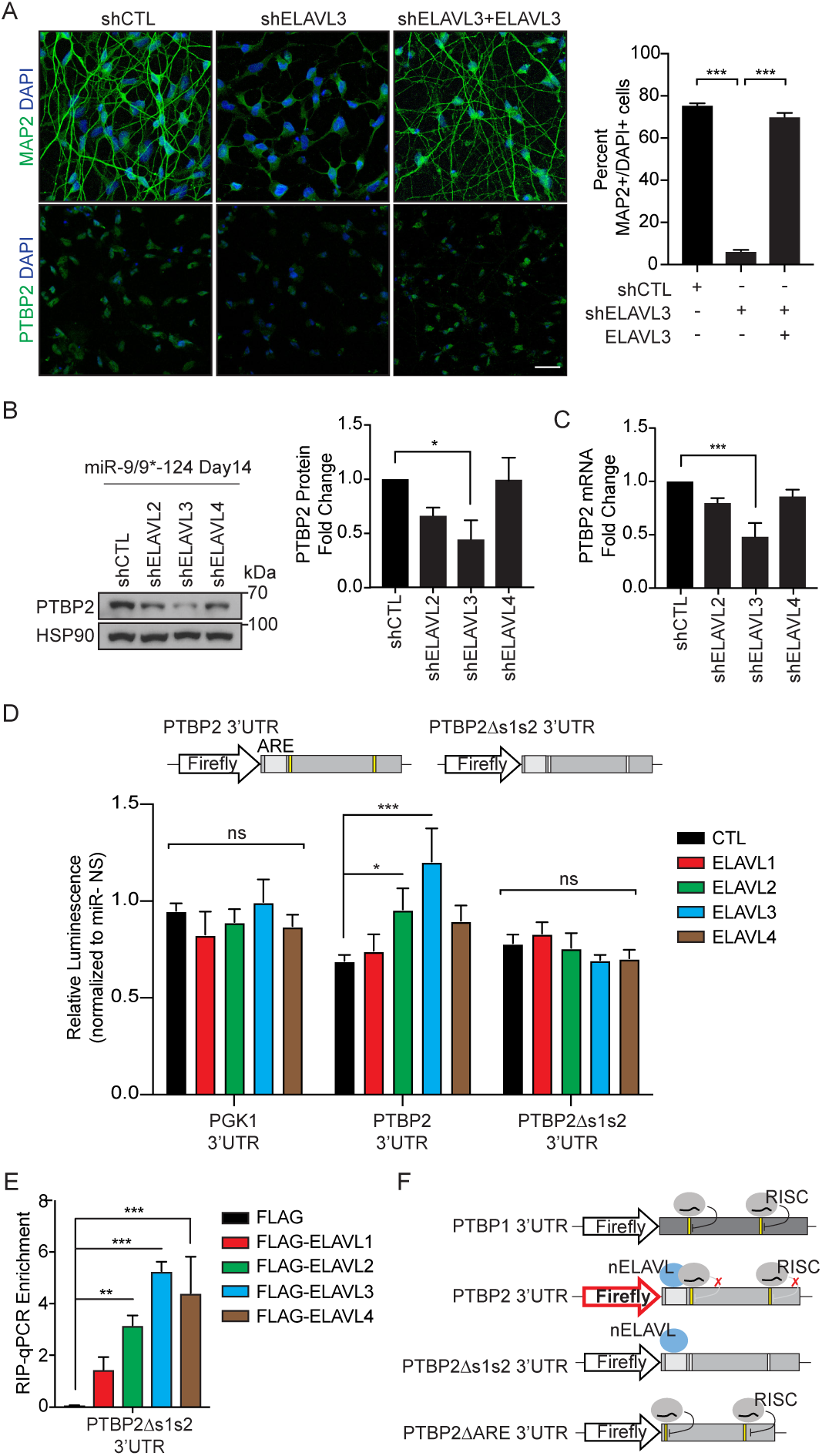
PTBP2 upregulation requires the synergism of nELAVL and miR-124. (A)Left, day 30 miNs with shRNA knockdown against CTL, ELAVL3, and ELAVL3 with ELAVL3 cDNA rescue. Cells were immunostained for MAP2 and PTBP2. Scale bar 50 = μm. Right, quantification of the percentage of MAP2-positive cells over the total number of DAPI-positive cells with two or more neurites (left). Data represented as mean ± SEM. One-way ANOVA followed by Tukey’s test (from left, *** P < 0.0001, *** P < 0.0001). shCTL n = MAP2 803/1060; shELAVL3 n = MAP2 67/1052; shELAVL3 + ELAVL3 n = MAP2 431/617. (B)Left, immunoblot analysis of PTBP2 in day14 miNs with knockdown against CTL, ELAVL2, ELAVL3, and ELAVL4. Right, quantification of the PTBP2 band intensity as a relative fold change compared to shCTL normalized to HSP90 in four independent experiments. Data are represented by mean ± SEM. One-way ANOVA followed by Dunnett’s test (* P = 0.0422). (C)The relative quantity of PTBP2 transcript upon individual nELAVL knockdown compared to CTL at day14 miNs determined by qPCR. Data are represented by mean ± SEM from 3 independent experiments. One-way ANOVA followed by Dunnett’s test (*** P = 0.0001). (D)Top, a diagram of luciferase constructs containing PTBP2 3’UTR with or without miR-124 sites (PTBP2Δs1s2). Bottom, luciferase assays with the addition of individual ELAVLs in PGK1 control, PTBP2, or PTBP2Δs1s2 3’UTR. Luminescence was measured 48 hrs after transfection and normalized miR-9/9*-124 to miR-NS control of each condition. Data are represented by mean ± SEM from at least three independent experiments.Two-way ANOVA followed by Dunnett’s test (* P = 0.0312, *** P = 0.0001). (E)RIP of individual FLAG-tagged ELAVLs in PTBP2Δs1s2 3’UTR 48 hrs after transfection. Enrichment normalized to input determined through qPCR against PTBP2 3’UTR. Data are represented by mean ± SEM from three independent experiments. Two-way ANOVA followed by Dunnett’s test (from left, PTBP2Δs1s2 3’UTR ** P = 0.0018, *** P = 0.0001, *** P = 0.0001). (F)A schematic summary of nELAVL and RISC activities on PTB 3’UTRs.

### Synergism between nELAVL and AGO requires miR-124 site in PTBP2 3’UTR

We further tested whether the miR-124 sites within *PTBP2* 3’UTR would also be critical for mediating the PTBP2 upregulation with ELAVL3. By mutating the miR-124 seed-match sequences within *PTBP2* 3’UTR (*PTBP2Δs1s2* 3’UTR), adding ELAVLs failed to enhance the luciferase activity over the control (CTL, black; relative luminescence ratio unchanged; *PTBP2Δs1s2* 3’UTR histograms) (Figure 5D). This result is in contrast to wild-type *PTBP2* 3’UTR where ELAVL3 can enhance luciferase activity in the presence of miR-124 target sites (Figures 4D, 5D). Interestingly, the lack of the increased luminescence with ELAVL3 addition was not due to the failure of ELAVL binding to *PTBP2Δs1s2* 3’UTR because qPCR analysis with nELAVL-RIP showed persistent binding of ELAVLs to *PTBP2Δs1s2* 3’UTR (Figure 5E). These results altogether suggest the requirement of both ARE and miR-124 sites in *PTBP2* 3’UTR for PTBP2 upregulation (Figure 5F).

### Neuronal genes downregulated upon loss of miR-124 and ELAVL3 in human neurons

To test whether the synergism of miR-124 and ELAVL3 for miRNA-mediated upregulation is unique to PTBP2 or a broader mechanism applicable to other genes beyond reprogrammed neurons, we performed loss-of-function studies on primary human neurons (HNs). With the same TuD-miR-124 and shELAVL3 constructs used in reprogrammed neurons, miR-124 and ELAVL3 were knockdown in HNs and processed for RNA-seq (Figures 6A-B and S6A). To identify potential targets upregulated by miR-124 and ELAVL3 in HNs, we focused on genes suppressed upon expression of TuD-miR-124 and shELAVL3 (KD, Log_2_FC ≤ −1; adj.P-value < 0.05) compared to CTL (Figure 6B). Many of these downregulated genes are neuronal-related, including *MAP2, PTBP2, SCN1A*, and *SEMA6A* (Figure 6B). Despite the downregulation of several neuronal transcripts, overall neuronal identity remains intact as expression of neuronal markers such as *RBFOX2, FMR1*, and *NEFL* remain similar between CTL and KD HNs (Figure S6B). Furthermore, we also do not observe an emergence of progenitor marker expression upon knockdown of both miR-124 and ELAVL3 (Figure S6B).

**Figure 6.**
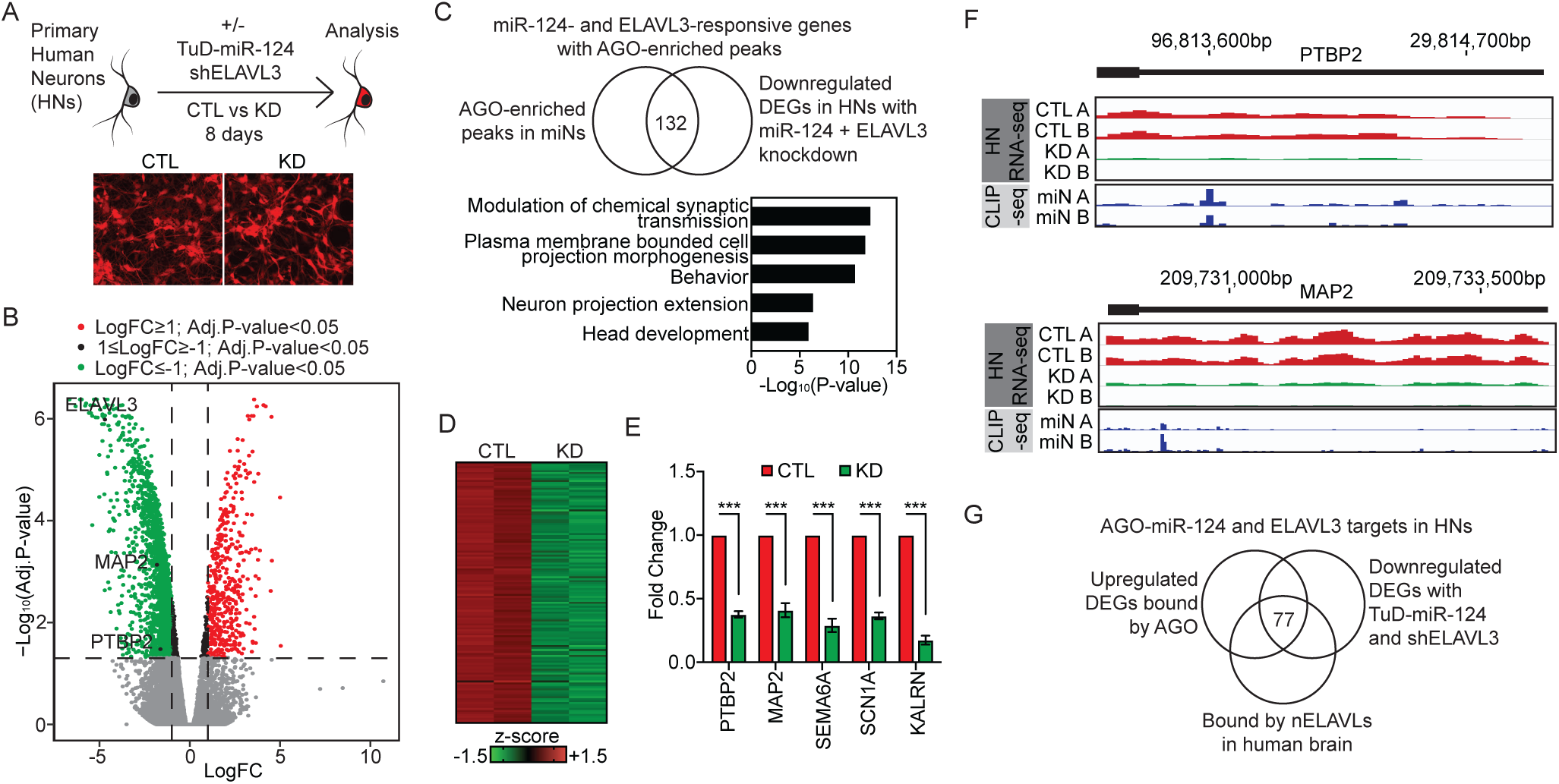
MiRNA-mediated upregulation of neuronal genes in primary human neurons. (A)Top, a schematic diagram of the experimental procedure using primary human neurons (HNs). Bottom, images of HNs marked by TurboRFP reporter in the non-specific miRNA or miR-124 tough decoy. (B)A volcano plot of differentially expressed genes between CTL and KD conditions with TuD-miR-124 and shELAVL3 treatment. A selection of downregulated genes are highlighted (ELAVL3, MAP2, and PTBP2). Red dots, HN mRNA expression Log2FC ≥ 1; adj.P-value < 0.05. Green dots, HN mRNA expression Log2FC ≤ −1; Adj.P-value < 0.05. (C)Top, by overlapping downregulated DEGs in (A) to previously identified upregulated AGO-enriched targets (Figure 1), 132 genes were identified to habor AGO peaks in miNs and are downregulated with miR-124 an ELAVL3 knockdown in HNs. Bottom, top biological GO terms associated with the 102 genes. (D)A heatmap of z-scores of the 132 genes identified in (B) from RNA-seq comparing KD and CTL in HNs. (E)RT-qPCR validation of a selection of the identified downregulated genes in HNs (C) that are found to be commonly targeted by both miNs and HNs (B). Data are represented in ± SEM from three independent experiments. Two-tailed unpaired t-test (all, *** P < 0.001). (F)Track views of HN RNA-seq tracks (top) and miN AGO HITS-CLIP tracks (bottom) for gene examples showing reduced expression upon knockdown of miR-124 and ELAVL3 in HNs (over CTL), and AGO-enriched peaks at the 3’UTR. (G)A venn diagram of AGO-miR-124 targets in HNs overlapped with nELAVL-bound targets in the human brain.

To ensure that these downregulated genes are miR-124 targets, we compared the downregulated DEGs from HNs to upregulated DEGs in day 20 miNs that also harbor AGO-enriched peaks (Figure 1). This comparison resulted in a set of 132 target genes associated with biological GO terms related to neuronal development and projection (Figure 6C, Table S3), further validating miR-124 as a positive regulator of the neuronal program. Examples of some these identified targets in HNs also validated in qPCR include *PTBP2, MAP2, SEMA6A, SCN1A*, and *KALRN* (Figures 6D-F). Together our data support the notion that miR-124-mediated upregulation of neuronal genes is not unique to PTBP2 in reprogramming context, but applicable to other neuronal genes in actual human neurons.

### MiR-124 and nELAVL interaction for other neuronal transcripts

Using existing nELAVL HITS-CLIP of the human brain (Scheckel et al., 2016), we performed a comparative analysis with our HNs dataset to examine if these upregulated neuronal transcripts are likely targets of nELAVLs. First, by overlapping i) upregulated neuronal DEGs bound by AGO HITS-CLIPs (Figure 1), ii) DEGs responsive to miR-124 tough decoy and *ELAVL3* shRNA in HNs (Figure 6), and iii) genes bound by nELAVLs in the human brain (Scheckel et al., 2016), we identified 77 genes, including *PTBP2, MAP2, SLCA48, KALRN, BCL7A*, and *SCN1A* enriched for neuronal biological terms generally involved in synaptic processes (Figures 6G and S7, Table S4). Based on these comparisons, similar to what we observed in miNs, miR-124 and ELAVL3 appear to collectively upregulate a set of neuronal genes that are likely critical for neuronal function in primary HNs.

### Neuronal properties affected by miR-124 and nELAVLs

As our AGO-HITS-CLIP and miR-124 knockdown data in miNs indicate that neuronal genes are preferentially targeted as cells acquire the neuronal fate (Figures 1 and 2), we examined the global transcriptome changes of our knockdown (KD) and control (CTL) conditions in HNs with LONGO analysis to assess changes in long gene expression (LGE), a measure of neuronal identity and maturation (Gabel et al., 2015; King et al., 2013; McCoy et al., 2018; McCoy and Fire, 2020; Sugino et al., 2014). Overall, knockdown of miR-124 and ELAVL3 (KD, green) in HNs resulted in reduced LGE compared to control (CTL, red), suggesting that both players are likely essential for the expression of long genes (Figure 7A).

**Figure 7.**
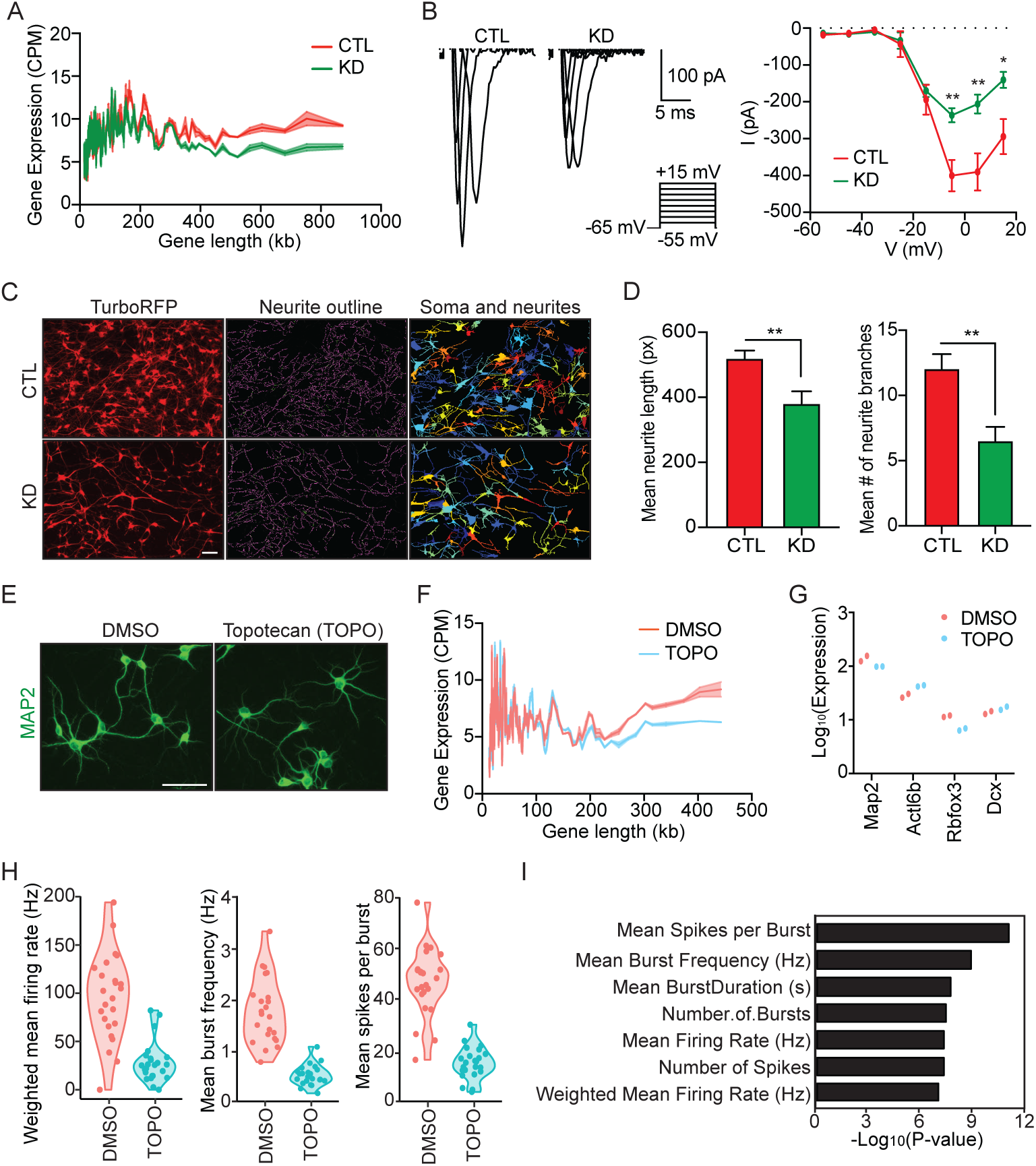
Long gene dysregulation results in altered neuronal properties. (A)LONGO plot showing reduced long gene expression upon knockdown of miR-124 and ELAVL3 (KD; green) compared to CTL (red). Lines show mean gene expression and ribbons show standard error of the mean (SEM). (B)Left, voltage-clamp traces of CTL and KD (TuD-miR-124 and shELAVL3) HNs. Right, average I-V curve of for all recorded CTL and KD HNs. Data are represented in ± SEM from seven recorded cells from each condition. Two-tailed unpaired t-test (from left, ** P = 0.00434; ** P < 0.00681; * P = 0.01233). (C)Representative images of CTL and KD HNs marked by TurboRFP reporter. Processed images by CellProfiler to identify neurites and associated cell soma. Scale bar = 100 μm. (D)Left, mean neurite length measurement of CTL and KD HNs. Data are represented in ± SEM from seven separate fields of view; CTL n = 768, KD n = 677. Two-tailed unpaired t-test (** P = 0.0092). Right, mean number of neurite branches in CTL and KD HNs. Data are represented in ± SEM from seven separate fields of view; CTL n = 768, KD n = 677. Two-tailed unpaired t-test (** P = 0.0038). (E)Representative images of primary rat neurons stained for MAP2 after treatment with DMSO control or Topotecan (TOPO). (F)LONGO plot showing reduced long gene expression upon treatment with TOPO (blue) compared to DMSO (red) in primary rat neurons. Lines show mean gene expression and ribbons show standard error of the mean (SEM). (G)Expression of a select unaffected neuronal markers between DMSO and TOPO treatments. (H)MEA readout for mean firing rate, burst frequency, and spikes per bursts of primary rat neurons treatment with DMSO or TOPO. (I)P-value of the various measured electrophysiological properties from MEA including those shown in (H).

As a number of identified neuronal genes targeted by AGO-miR-124 and ELAVL3 in HNs are implicated in neuronal function and morphology such as *SCN1A, SLC4A8, ANK3*, and *MAP2*, reflective of reduced overall LGE (Figures 7A and S7A, Table S4), we examined a few neuronal properties in CTL and KD HNs. We first performed electrophysiology and found KD HNs exhibited reduced inward sodium current as compared to CTL HNs (Figures 7B and S6C-D) which we reasoned to be attributed to a number of downregulated channel genes in KD HNs. However, other electrical properties such as resting membrane potential and action potential firing appear similar between CTL and KD HNs (Figures S6D-E). In addition, as we observed reduced neurite complexity in our KD cells compared to CTL HNs, we sought to measure features such as average neurite length and average number of neurite branches between the two conditions. Overall, we found that KD HNs not only have shorter average neurite length per cell, but also have fewer branches per cell compared to CTL (Figures 7C-E).

Reduced LGE in primary HNs with both miR-124 and ELAVL3 knockdown suggests that LGE is a transcriptomic phenotype reflective of altered neuronal features observed and measured in HNs. To independently validate the correlation between LGE and neuronal maturity, we also treated primary rat neurons with Topotecan (TOPO), an inhibitor of Topoisomerase I known to reduce LGE in neurons (King et al., 2013; Mabb et al., 2016, 2014). Primary rat neurons treated with TOPO (blue) were found to display a reduction in LGE when compared to the control condition (DMSO, red) (Figures 7E-F) while the TOPO-treated cells still maintained the expression of other neuronal markers such as *Map2, Actl6b, Rbfox3* and *Dcx* (Figure 7G). To further assess the consequences of LGE reduction in neurons, we measured electrophysiological properties using a microelectrode array (MEA). TOPO-treated neurons exhibited altered electrophysiological properties compared to DMSO control, including reduced mean firing rate, mean burst frequency, and number of spikes per burst (Figures 7H-I), demonstrating that LGE is a transcriptomic feature related to the functional maturity of neurons. Therefore, our results support the notion that miR-124, in synergy with ELAVL3, promote neuronal maturity by positively regulating their target genes, as evidenced by their effect on LGE in human neurons. Based on PTBP2, we delineated a mechanism on how miR-124 and ELAVL3 can promote the expression of their targets and we anticipate that some of the additional identified long genes important for neuronal differentiation and function are likely upregulated in a similar manner.

## Discussion

In the present study, we uncovered the role of miR-124 in the upregulation of genes associated with neuronal differentiation and function during the neuronal conversion of human fibroblasts. This finding provides insights into the function of miRNAs in addition to their canonical role as a repressor of downstream target genes. Of the bound AGO transcripts are *bona fida* miR-124 target genes in which the 3’UTRs can be repressed in a non-neuronal context, but reverses upon neuronal induction. By examining the mutually exclusive regulation of *PTBP1* and *PTBP2*, we reveal how miR-124 plays a bifunctional role depending on the sequence composition at the 3’UTR, the availability of neuronal ELAVLs, and the interaction with AGOs to mediate the switching of PTB homolog expression during the neuronal conversion of HAFs. Although we focus specifically on the interplay between miR-124 and ELAVL3 for PTBP2 upregulation, future studies should also examine if similar mechanism is used to promote other identified neuronal transcripts or if other RBPs can also synergize with AGOs for such target gene regulation. For example, FXR1, an RBP, has been shown in previous studies to interact with AGOs to facilitate gene expression in non-neuronal cells (Truesdell et al., 2012; Vasudevan et al., 2007; Vasudevan and Steitz, 2007).

The selective role of ELAVL3, and not ubiquitous ELAVL1, in mediating PTBP2 induction in neurons highlights the functional specificity of ELAVL family members in neurons. Like ELAVL3, ELAVL1 binds AREs and has been shown to interact with RISC components (Kim et al., 2009; Vasudevan and Steitz, 2007). Although different studies reveal opposing consequence of RISC and ELAVL family interaction on target genes, downstream functional output of AGO-ELAVL likely depends on not only the concurrent availability of RBP and target transcript, but also the 3’UTR sequence. Our results define the functional specificity inherent in ELAVL3 that cannot be replaced by ELAVL1, especially for regulating PTBP2 expression with miR-124 in neurons. We found that the specificity is, at least in part, driven by the hinge region inferred by the region-swapping experiments between ELAVL1 and ELAVL3. As there is no known ARE recognition unique to each ELAVL family member, it is likely that interactors associating with the hinge region may be regulating ELAVL specificity and targeting (Fujiwara et al., 2012; Hinman et al., 2013).

To investigate if the synergism of miR-124 and nELAVLs for transcript stabilization and activation can be generalized to other neuronal transcripts beyond *PTBP2* outside the conversion system, we also knockdown miR-124 and ELAVL3 in primary human neurons. Our results indicate that several neuronal transcripts are induced by miR-124 and ELAVL3 that are critical for neuronal program. Furthermore, by examining existing nELAVL HITS-CLIP datasets in the human brain (Scheckel et al., 2016), we uncovered that miR-124- and nELAVL-mediated upregulation of target transcripts may not be a unique occurrence to *PTBP2*, but likely an overlooked mechanism that maintains gene expression in neurons. Interestingly, as we identified numerous neuronal genes to be targets of miR-124, such as *MAP2, CAMK1D*, and *SEMA6A*, we argue that miR-124 may be critical for enhancing the overall neuronal program as measured by the reduced sodium current, and neurite length and branches through its regulation on long genes. By chemically inhibiting LGE in rat neurons, mimicking the transcriptomic phenotype observed in HNs with reduce miR-124 and ELAVL3 activity, we observed a variety of altered electrical properties. This finding also indicates that miR-9/9*-124 is highly neurogenic as these miRNAs not only allow for the conversion of HAFs into neurons (Abernathy et al. 2017; Yoo et al. 2011), but also enhanced the expression of neuronal markers, such as *MAP2*, when overexpressed during neuronal differentiation of human pluripotent stem cell-derived neurons (Ishikawa et al., 2020; Sun et al., 2016). Although further tests are required to see if all the identified transcripts targeted by both miR-124 and nELAVLs share the same mechanism as *PTBP2*, our loss-of-function results from in both reprogramming and primary human neuron systems, and its effect on other neuronal genes lend support to the general role of miR-124 as a positive regulator of select target genes in neurons. Future experiments taking a closer look at RNA structure, motif proximity with RBPs, and additional interacting proteins will provide further mechanistic insights to how a single miRNA can simultaneously repress and activate different transcripts in a cell context-dependent manner.

Our study offers insights into the bifunctional mode of miR-124 conferring miRNAs as a reprogramming effector that can contribute to the neuronal program by upregulating specific neuronal genes. As miRNAs have been typically checked for their targets in non-neuronal cell lines, it is plausible that the dual-mode of miRNAs may not be unique to miR-124 in neurons, but instead utilized and altered in other cellular contexts with the help of cell type-specific RBPs. The governance of miRNA activity by the sequences within 3’UTR highlights the role of cell type-specific RBPs as core regulators of gene expression.

## Materials and Methods

### Cell culture

Primary human fibroblasts used in this study was from a 22-year-old female (GM02171, NIGMS Coriell Institute for Medical Research) while human neonatal fibroblasts (ScienCell, 2310) was used exclusively for the HITS-CLIP experiment. Fibroblasts were maintained in high glucose Dulbecco’s Modified Eagle Medium (Gibco, 11960044) containing 10% FBS (Gibco, 10437028), MEM non-essential amino acids (Gibco, 11140050), sodium pyruvate (Gibco, 11360070), GlutaMAX (Gibco, 35050061), HEPES (Gibco, 15630080), penicillin-streptomycin (Gibco, 15130122), and 2-mercaptoethanol (Gibco, 21985023) at 37°C. Primary human neurons were obtained commercially (ScienCell, 1520) with gender and age of the source undisclosed Human neurons were maintained in neuronal media (NM; ScienCell, 1521) at 37°C. E18 rat cortex (BrainBits®, FSDECX1M) were grown in STEMdiff™Neural Induction Medium (STEMCELL Technologies, 05835).

Lenti-X 293T (Clontech, 632180) cells were maintained in high glucose Dulbecco’s Modified Eagle Medium (Gibco, 11960044) containing 10% FBS (Gibco, 10437028), MEM non-essential amino acids (Gibco, 11140050), sodium pyruvate (Gibco, 11360070), GlutaMAX (Gibco, 35050061), HEPES (Gibco, 15630080), penicillin-streptomycin (Gibco, 15130122), and 2-mercaptoethanol (Gibco, 21985023) at 37°C.

### MiR-9/9*-124-mediated neuronal conversion

To initiate reprogramming, doxycycline-inducible pT-BclXL-miR-9/9*-124 (Addgene, 60857) and reverse tetracycline-controlled transactivator rtTA (Addgene, 66810) lentivirus with 8 ug/mL polybrene (Sigma, H9268) was added to a plate of confluent fibroblasts and spinfected at 37°C for 30 min at 1,000xG. Full media change with 1 μg/mL doxycycline (DOX; Sigma-Aldrich, D9891) occurred the following day. Two days following transduction, cells underwent another media change supplemented with DOX and respective antibiotics (Puromycin, Life Technologies, A11138-03; Blasticidin S HCl, Life Technologies, A11139-03). Five days after transduction, cells were plated onto poly-l-ornithine (Sigma-Aldrich, P4957), fibronectin (Sigma-Aldrich, F4759), and laminin (Sigma-Aldrich, L2020) coated coverslips or onto 10cm^2^ Primaria plates (Corning, 353803) followed by full media switch to neuronal media (NM; ScienCell, 1521) the next day supplemented with 200 μM dibutyl-cyclic AMP (cAMP; Sigma-Aldrich, D0627), 1 mM valproic acid (VPA; Sigma Aldrich, P4543), 10 ng/mL human BDNF (PeproTech, 450-02), 10 ng/mL human NT-3 (Peprotech, 450-03), 1 μM retinoic acid (RA; Sigma-Aldrich, R2625), RevitaCell supplement (Gibco, A2644501), and antibiotics. DOX was supplemented every 2 days while half media changes occurred every 4 days until day 30.

### Plasmids and cloning

For luciferase assay, full length 3’UTR of target transcripts were cloned and ligated into pmirGLO vector. For mutagenizing miR-124 target sites, QuikChange XL site-directed mutagenesis kit (Agilent, 200516) was used according to the manufacturer’s protocol. Using the same UTR sequences, 3’UTR was attached immediately downstream of a destablized EGFP reporter and subcloned into lentiviral vector. Sequences for shRNAs were synthesized through Integrated DNA Technologies, annealed, and ligated into the pLKO.1 vector (Addgene, 8453 or 26655). Overexpression vectors of either lentiviral (N106 or N174) or mammalian expression (pcDNA3.1+, Invtrogen, V79020) was cut using NotI cut site (NEB, R0189) for insert ligation. Tough decoy for miR-124 was synthesized (GeneScript) and subcloned into pLemir vector using MluI (NEB, R0198) and NotI sites.

### Lentivirus production

Lentivirus was produced as previously described (Richner et al., 2015). Briefly, 1.5 μg pMD2.G, 4.5 μg psPAX2, 6 μg of plasmid in lentiviral backbone, 600 μl Opti-MEM (Life Technologies, 31985) and 48 μl of 2 mg/mL polyethyleneimine (PEI; Polysciences, 24765) were mixed and transfected into Lenti-X 293T (Clontech, 632180) plated at 6×10^6^ cells per 10 cm^2^ dish. Media was changed the following day, and viral supernatant was collected, filtered and spun at 70,000xG for 2 hr at 4°C two days later. The viral pellet collected per 10 cm^2^ dish was resuspended in 1 mL PBS.

### Immunostaining analysis

Cells were fixed with 4% paraformaldehyde (PFA; Electron Microscopy Sciences, 15710) for 20 mins at room temperature (RT) followed by three washes with PBS. Cells were permeabilized and blocked in 0.3% TritonX-100, 2% normal goat serum (NGS; Jackson ImmunoResearch Laboratories, 005-000-121) and 5% bovine serum albumin (BSA; Sigma-Aldrich, A7906) in PBS for 1 hr at RT prior to incubation with primary antibodies overnight at 4°C. After three washes with PBS, cells were incubated with respective secondary antibodies for 1 hr at RT. Coverslips were mounted onto coverslides with ProLong Gold antifade reagent (Invitrogen, P36934) for imaging using Leica SP5X white light laser confocal system with Leica Application Suite (LAS) Advanced Fluorescence. See Supplemental Table S4 for a list of antibodies used.

### SYTOX assay

SYTOX assay was performed as previously described (Victor et al., 2018). Briefly, 0.1 μM SYTOX gene nucleic acid stain (Invitrogen, S7020) and 1 μl/mL of Hoeschst 33342 (Thermo Scientific, 66249) were added into cell medium. Samples were incubated for at least 15 mins in 37°C prior to imaging. Images were taken using Leica DMI 4000B inverted microscope with Leica Application Suite (LAS) Advanced Fluorescence.

### Luciferase assay

HEK 293 cells plated in 96-well plate were transfected with 100 ng of pSilencer-miRNA, 100 ng of pmirGLO containing 3’UTR of interest, and PEI (Polysciences, 24765) with Opti-MEM (Life Technologies, 31985). Forty-eight hours after transfection, luciferase activity was assayed using Dual-Glo luciferase assay system (Promega, E2920) according to the manufacturer’s protocol using Synergy H1 Hybrid plate reader (BioTek). Luciferase activity was obtained by normalizing firefly luminescence to renilla luminescence (luciferase activity = firefly/renilla) followed by normalizing to respective pSilencer-miR-NS control.

### Flow cytometry

Destabilized EGFP reporter with or without 3’UTR of interest was transduced into HEK 293 cells to establish a stable reporter containing cell line with Blasticidin S HCl (Life Technologies, A11139-03) selection. The day following transduction with miR-9/9*-124 lentivirus, media was changed with the addition of DOX (Sigma-Aldrich, D9891). At day 3, media change was supplemented with DOX and puromycin (Life Technologies, A11138-03). DOX was supplemented every 2 days following transduction. At day 10, cells were imaged, and collected for flow cytometry. Briefly, cells were collected in PBS and incubated with propidium iodide (PI; Sigma-Aldrich, P4861) on ice until ready. Using FACSCalibur (BD Biosciences), all PI-negative cell population was obtained for the gating of GFP-negative and -positive cell population.

### Quantitative reverse transcription PCR

Total RNA of cells was extracted using TRIzol Reagent (Invitrogen, 15596026). Reverse transcription was performed using SuperScript III first strand synthesis system for RT-PCR (Invitrogen, 18080-051) according to the manufacturer’s protocol from fibroblasts, reprogrammed neurons, and human brain total RNA (Invitrogen, AM7962). Quantitative PCR was performed using SYBR Green PCR master mix (Applied Biosystems, 4309155) and StepOnePlus Real-Time PCR system (Applied Biosystems, 4376600) according to the manufacturer’s protocol against target genes.

### HITS-CLIP

AGO HITS-CLIP was performed on cells after 2 weeks into reprogramming of miR-NS or miR-9/9*-124-expressing neonatal fibroblasts (ScienCell, 2310) at day 14 and day 21. Cells were harvested, UV-crosslinked, lysed, and processed according to Moore et al. 2014. Briefly, cross-linked cells were lysed and treated with RQ1 DNase (Promega, M6101) and RNaseA (Thermo). Complex containing AGO-miRNA-mRNA were immunoprecipitated overnight at 4°C with pan-AGO antibody. The immunoprecipitated complex was radio-labelled and extracted after running on NuPAGE gel (Thermo). RNA from the 130kDa band was extracted for sequencing using TruSeq Small RNA Library Preparation Kits (San Diego, CA). Samples were sequences using Illumina HiSeq 2500 platform at the Genome Technology Access Center (GTAC) at Washington University School of Medicine, St. Louis.

### MiRNA Tough Decoy RNA-seq

Total RNA was extracted from day 20 cells expressing miR-9/9*-124 + TuD-miR-NS, and miR-9/9*-124 + TuD-miR-124 using TRIzol Reagent (Invitrogen, 15596026) in combination with RNeasy micro kit (Qiagen, 74004). RNA quality (RIN ≥ 9.6) was determined with 2100 Bioanalyzer (Agilent) and samples underwent low input Takara-Clontech SMARTer kit (Takara, 639490) library preparation. Samples were sequenced using NovaSeq S4 and processed at the Genome Technology Access Center (GTAC) at Washington University School of Medicine, St. Louis. For human neurons, total RNA was extracted from HNs after 8 days of tough decoy and shRNA treatment using TRIzol Reagent (Invitrogen, 15596026) in combination with RNeasy micro kit (Qiagen, 74004). RNA quality (RIN ≥ 8.4) was determined with 2100 Bioanalyzer (Agilent) and samples underwent TruSeq Stranded total RNA sequencing kit library preparation. Samples were sequenced using NovaSeq6000 through DNA Link (San Diego, CA).

### Human Clariom D Microarray

For Human Clariom D Array (Affymetrix), total RNA was extracted from day 14 cells expressing miR-NS, miR-9/9*-124 + shCTL, and miR-9/9*-124 + shPTBP2 using TRIzol Reagent in combination with RNeasy mini kit (Qiagen, 74104). RNA quality (RIN > 9.6) was determined with 2100 Bioanalyzer and samples (biological duplicates each) underwent amplification and hybridization according to manufacturer’s protocol by GTAC at Washington University School of Medicine, St. Louis.

### Immunoblot analysis

Cells were lysed with sonication (Diagenode, UCD-200) in RIPA buffer (Thermo Scientific, 89900) supplemented with protease inhibitor cocktail tablet (Roche, 04693132001). Protein concentration of cleared lysate was measured using Pierce BCA protein assay kit (Thermo Scientific, 23227) and read with Synergy H1 Hybrid plate reader (BioTek). Lysate and sample buffer (Life Technologies, NP0008) were boiled, separated with Bis-Tris gels, and transferred to nitrocellulose membrane (GE Healthcare Life Sciences, 10600006). Membrane was blocked with 5% milk for 1 hr at RT and incubated with primary antibody overnight at 4°C. After three washes of TBST (1X TBS and 0.1% Tween-20), the membrane was incubated with respective horseradish peroxidase-conjugated antibody for 1 hr at RT followed by three washes with TBST. Blots were developed with ECL system (Thermo Scientific, 34580) and imaged or developed onto film. See Supplemental Table S3 for a list of antibodies used.

### Immunoprecipitation analysis

Cells were lysed with sonication (Diagenode, UCD-200) in IP buffer (20 mM HEPES, 150 mM NaCl, 10% glycerol, 5 mM EDTA, 1% Triton X-100) supplemented with protease inhibitor cocktail tablet (Roche, 04693132001). Cleared lysate was incubated with anti-FLAG M2 magnetic beads (Sigma-Aldrich, M8823) overnight with rotation at 4°C. The beads were washed three times with IP buffer and bound proteins were boiled and eluted with sample buffer (Life Technologies, NP0008), separated with Bis-Tris gels, and immunoblotted.

### RNA-IP

Cells were lysed with sonication (Diagenode, UCD-200) in IP buffer (20 mM HEPES, 150 mM NaCl, 10% glycerol, 5 mM EDTA, 1% Triton X-100) supplemented with protease inhibitor cocktail tablet (Roche, 04693132001). Cleared lysate was incubated with anti-FLAG magnetic beads (Sigma-Aldrich, M8823) overnight with rotation at 4°C beads. The beads were washed three times with IP buffer and resuspended in 90 ul of IP buffer with proteinase K (NEB, P8107S) for 30 mins at 37°C. To extract RNA, 1 mL of TRIzol Reagent (Invitrogen, 15596026) was added to the bead slurry. Final RNA was DNase I (Invitrogen, 18068015) treated prior to RT-qPCR.

### Electrophysiology

Whole-cell patch-clamp recordings were performed as previously described (Victor et al., 2018). Briefly, HNs (ScienCell, 1521) were recording within 8 to 10 days after tough decoy and shRNA transduction. Recordings were acquired using pCLAMP 10 software, multipliclamp 700B amplifier, and Digidata 1550 digitizer (Molecular Devices, CA). Glass electrode pipettes were pulled from borosilicate glass (1B120F-4, World Precision) to obtain pipette resistance ranging from 5 – 8 MΩ using next generation micropipette puller (P-1000, Sutter Instrument). External solution is consist of 140 mM NaCl, 3 mM KCl, 10 mM Glucose, 10 mM HEPES, 2 mM CaCl_2_ and 1 mM MgCl_2_, and internal solution is consist of 130 mM K-Gluconate, 4 mM NaCl, 2 mM MgCl_2_, 1 mM EGTA, and 10 mM HEPES (adjusted to pH 7.25 with KOH) were used for recording. For all recordings, the membrane potentials were held at −65 mV.

### Microelectrode array

E18 rat cortical neurons (BrainBits, FSDECX1M) were plated onto a 24-well cell culture microelectrode array plate (Axion Biosystems, M384-tMEA-24W). Prior to plating, 1 ml of sterile water and 10 μl of PEI (Sigma-Aldrich, 03880) at 0.5% diluted in 0.1 M HEPES pH 8.0 (Caymen Chemical Company, 700014) was added to the center of each well. After incubating overnight at 37 °C, each well was washed 4x with sterile water, and allowed to dry for 15 minutes. 10 μl of Laminin (Sigma-Aldrich, L2020) at 10 μg/mL diluted in cold DMEM/F-12 (ThermoFisher, 11320082) was added to each well and incubated at 37 °C for 2 hours. Laminin was aspirated prior to plating neurons. 24 hours after plating, neurons were cultured in BrainPhys Neuronal Medium and SM1 supplement (STEMCELL Technologies, 05792) with either Topotecan (Sigma-Aldrich, T2705) at a final concentration of 300 nM in DMSO (0.05% DMSO final concentration) or DMSO control for one week before recording electrophysiological activity using a Maestro MEA plate reader (Axion Biosystems).

### GO enrichment analysis

Gene ontology analyses were performed using Metascape (Tripathi et al., 2015) with minimum overlap of 3, P-value cutoff of 0.01, and minimum enrichment of 1.5. Entire gene list was used for each GO analysis.

### RNAhybrid miRNA-target duplex analysis

To predict the presence of miR-124-3p binding sites at HITS-CLIP peaks, peak sequences were extracted by on the genomic coordinates and processed through RNAhybrid (Rehmsmeier, 2004) against miR-124-3p miRNA sequence with at least a free energy threshold of −20 kcal/mol.

### RBP binding analysis

RNA-binding protein prediction database/software, RBPDB (Cook et al., 2011), RBPmap (Paz et al., 2014), and beRBP (Yu et al., 2018) for *PTBP2* 3’UTR sequence were used. Default criteria were selected for all three software for non-bias prediction of any human RBP motifs.

### Neurite length measurement

Images of HNs marked by TurboRFP were processed through CellProfiler 3.1.9 (McQuin et al., 2018). Briefly, neurite feature was enhanced prior to the identification of primary object or soma between 25 – 100 pixel unit in diameter, followed by the identification of secondary object based on neurite feature. A morphological skeleton was made based on overlaying the identified objects and mean neurite lengths and branches were then measured using measure object skeleton module.

### LONGO analysis

LGE for human and rat neurons were determined by through the LONGO platform (McCoy et al., 2018). LONGO analysis output of gene expression (CPM) over gene length was used to generate the LONGO plot.

### Data Analyses

AGO HITS-CLIP reads for miR-NS and miR-9/9*-124-expressing cells at least two weeks into reprogramming were trimmed and aligned to human genome hg38 using STAR with default parameters. Differential peaks between miR-9/9*-124 and miR-NS conditions were detected using MACS (Feng et al., 2012; Zhang et al., 2008) with the following criteria: mfold bound of 5 – 50, fragment size of 100, and FDR > 0.05. HITS-CLIP datasets will be publicly available.

RNA-seq for day 20 miNs expressing either CTL TuD-miR-NS or TuD-miR-124 was analyzed by GTAC’s RNA-seq pipeline. Briefly, reads were aligned to hg38 with STAR and processed through EdgeR (Robinson et al., 2010) to obtain differentially expressed genes with adj. p-value of < 0.01. RNA-seq dataset consisting of TuD-miR-NS and TuD-miR-124 at day 20 will be publicly available.

RNA-seq for HNs treated with either CTL or tough decoy against miR-124 and shRNA against ELAVL3 (KD) was aligned through Partek^®^ Flow^®^ software (Partek Inc., 2020) to generate gene counts. Briefly, reads were aligned to hg38 with STAR and processed through EdgeR (Robinson et al., 2010) to obtain differentially expressed genes with adj. p-value of < 0.05 and Log_2_FC of ≤ −1 and ≥ 1. RNA-seq dataset consisting of CTL and KD treatments from HNs will be publicly available.

RNA-seq for primary rat neurons treated with either DMSO or Topotecan (TOPO) were aligned to the Ensembl release 76 top-level assembly with STAR. Transcript counts were produced by Sailfish version 0.6.3. All gene-level and transcript counts were then imported into the R/Bioconductor package EdgeR and TMM normalization size factors were calculated to adjust samples for differences in library size. RNA-seq dataset consisting of DMSO and TOPO treatments from rat neurons will be publicly available.

Human Clariom D Array data was analyzed using manufacturer’s software, Expression Console followed by Transcriptome Analysis Console (TAC). For gene level analysis, genes comparing miR-9/9*-124 vs miR-NS linear fold change of ≥ 1.5 and ANOVA P < 0.05 were considered to upregulated in neuronal conditions. For splicing analysis, splicing events with linear splicing index ≤ −2 and ≥ 2, ANOVA P < 0.05 were considered significant splicing events. Human Clariom D array dataset consisting of miR-NS, miR-9/9*-124 with shCTL, and miR-9/9*-124 with shPTBP2 at day 14 will be publicly available.

## Author Contributions

Y.L.L. conducted experiments shown in Figures 2-7 and associated supplementary Figures 1, 3-Y.J.L. conducted experiments shown in Figure 1 and supplementary Figure 2. M.J.M. conducted experiments shown in Figure 7. A.S.Y. supervised the study. Y.L.L. and A.S.Y. wrote the manuscript.

## Acknowledgements

We thank the GTAC at Washington University for the sequencing service and support, and flow Cytometry Core at Washington University for equipment use. We thank We thank P. Gontarz, for the bioinformatics analysis of AGO HITS-CLIP, D. Annamalai and A. H. Kim for the help with tough decoy designs, M. Victor for the help with electrophysiology, and K. Cates and L. Capano for the helpful suggestions with the manuscript. Y.L.L. is supported by the LIFENAD Fellowship. M.J.M. was supported by an IPNG fellowship (T32GM081739; Barch, PI). This study was supported through awards and funds to A.S.Y by the Andrew B. and Virginia C. Craig Faculty Fellowship endowment, NIH Director’s Innovator Award (DP2NS083372), Presidential Early Career Award for Scientists and Engineers (PECASE), and NIH (RF1AG056296 and R01NS107488).

## Declaration of Interests

The authors declare no competing interest.

## Notes

### Competing Interest Statement

The authors have declared no competing interest.

## References

Abernathy DG, Kim WK, McCoy MJ, Lake AM, Ouwenga R, Lee SW, Xing X, Li D, Lee HJ, Heuckeroth RO, Dougherty JD, Wang T, Yoo AS. 2017a. MicroRNAs Induce a Permissive Chromatin Environment that Enables Neuronal Subtype-Specific Reprogramming of Adult Human Fibroblasts. Cell Stem Cell 21:332-348.e9. doi:10.1016/j.stem.2017.08.002

Bak RO, Hollensen AK, Primo MN, Sorensen CD, Mikkelsen JG. 2013. Potent microRNA suppression by RNA Pol II-transcribed “Tough Decoy” inhibitors. RNA 19:280–293. doi:10.1261/rna.034850.112

Ballas N, Grunseich C, Lu DD, Speh JC, Mandel G. 2005. REST and Its Corepressors Mediate Plasticity of Neuronal Gene Chromatin throughout Neurogenesis. Cell 121:645–657. doi:10.1016/j.cell.2005.03.013

Boutz PL, Stoilov P, Li Q, Lin C-H, Chawla G, Ostrow K, Shiue L, Ares M, Black DL. 2007. A post-transcriptional regulatory switch in polypyrimidine tract-binding proteins reprograms alternative splicing in developing neurons. Genes & Development 21:1636–1652. doi:10.1101/gad.1558107

Cook KB, Kazan H, Zuberi K, Morris Q, Hughes TR. 2011. RBPDB: a database of RNA-binding specificities. Nucleic Acids Research 39:D301–D308. doi:10.1093/nar/gkq1069

Feng J, Liu T, Qin B, Zhang Y, Liu XS. 2012. Identifying ChIP-seq enrichment using MACS. Nat Protoc 7:1728–1740. doi:10.1038/nprot.2012.101

Fujiwara T, Fukao A, Sasano Y, Matsuzaki H, Kikkawa U, Imataka H, Inoue K, Endo S, Sonenberg N, Thoma C, Sakamoto H. 2012. Functional and direct interaction between the RNA binding protein HuD and active Akt1. Nucleic Acids Research 40:1944–1953. doi:10.1093/nar/gkr979

Gabel HW, Kinde B, Stroud H, Gilbert CS, Harmin DA, Kastan NR, Hemberg M, Ebert DH, Greenberg ME. 2015. Disruption of DNA-methylation-dependent long gene repression in Rett syndrome. Nature 522:89–93. doi:10.1038/nature14319

Haraguchi T, Ozaki Y, Iba H. 2009. Vectors expressing efficient RNA decoys achieve the long-term suppression of specific microRNA activity in mammalian cells. Nucleic Acids Research 37:e43–e43. doi:10.1093/nar/gkp040

Hinman MN, Zhou H-L, Sharma A, Lou H. 2013. All three RNA recognition motifs and the hinge region of HuC play distinct roles in the regulation of alternative splicing. Nucleic Acids Research 41:5049–5061. doi:10.1093/nar/gkt166

Iadevaia V, Gerber A. 2015. Combinatorial Control of mRNA Fates by RNA-Binding Proteins and Non-Coding RNAs. Biomolecules 5:2207–2222. doi:10.3390/biom5042207

Ince-Dunn G, Okano HJ, Jensen KB, Park W-Y, Zhong R, Ule J, Mele A, Fak JJ, Yang C, Zhang C, Yoo J, Herre M, Okano H, Noebels JL, Darnell RB. 2012. Neuronal Elav-like (Hu) Proteins Regulate RNA Splicing and Abundance to Control Glutamate Levels and Neuronal Excitability. Neuron 75:1067–1080. doi:10.1016/j.neuron.2012.07.009

Ishikawa M, Aoyama T, Shibata S, Sone T, Miyoshi H, Watanabe H, Nakamura M, Morota S, Uchino H, Yoo AS, Okano H. 2020. miRNA-Based Rapid Differentiation of Purified Neurons from hPSCs Advancestowards Quick Screening for Neuronal Disease Phenotypes In Vitro. Cells 9:532. doi:10.3390/cells9030532

Jiang P, Coller H. 2012. Functional Interactions Between microRNAs and RNA Binding Proteins. MicroRNA e 1:70–79. doi:10.2174/2211536611201010070

Kim HH, Kuwano Y, Srikantan S, Lee EK, Martindale JL, Gorospe M. 2009. HuR recruits let-7/RISC to repress c-Myc expression. Genes & Development 23:1743–1748. doi:10.1101/gad.1812509

King IF, Yandava CN, Mabb AM, Hsiao JS, Huang H-S, Pearson BL, Calabrese JM, Starmer J, Parker JS, Magnuson T, Chamberlain SJ, Philpot BD, Zylka MJ. 2013. Topoisomerases facilitate transcription of long genes linked to autism. Nature 501:58–62. doi:10.1038/nature12504

Lee SW, Oh YM, Lu Y-L, Kim WK, Yoo AS. 2018. MicroRNAs Overcome Cell Fate Barrier by Reducing EZH2-Controlled REST Stability during Neuronal Conversion of Human Adult Fibroblasts. Developmental Cell 46:73-84.e7. doi:10.1016/j.devcel.2018.06.007

Lessard J, Wu JI, Ranish JA, Wan M, Winslow MM, Staahl BT, Wu H, Aebersold R, Graef IA, Crabtree GR. 2007. An Essential Switch in Subunit Composition of a Chromatin Remodeling Complex during Neural Development. Neuron 55:201–215. doi:10.1016/j.neuron.2007.06.019

Li Q, Zheng S, Han A, Lin C-H, Stoilov P, Fu X-D, Black DL. 2014. The splicing regulator PTBP2 controls a program of embryonic splicing required for neuronal maturation. eLife 3. doi:10.7554/eLife.01201

Licatalosi DD, Yano M, Fak JJ, Mele A, Grabinski SE, Zhang C, Darnell RB. 2012. Ptbp2 represses adult-specific splicing to regulate the generation of neuronal precursors in the embryonic brain. Genes & Development 26:1626–1642. doi:10.1101/gad.191338.112

Lin K, Zhang S, Shi Q, Zhu M, Gao L, Xia W, Geng B, Zheng Z, Xu EY. 2018. Essential requirement of mammalian *Pumilio* family in embryonic development. MBoC 29:2922–2932. doi:10.1091/mbc.E18-06-0369

Lu J, Ji H, Tang H, Xu Z. 2018. microRNA-124a suppresses PHF19 over-expression, EZH2 hyper-activation, and aberrant cell proliferation in human glioma. Biochemical and Biophysical Research Communications 503:1610–1617. doi:10.1016/j.bbrc.2018.07.089

Mabb AM, Kullmann PHM, Twomey MA, Miriyala J, Philpot BD, Zylka MJ. 2014. Topoisomerase 1 inhibition reversibly impairs synaptic function. Proc Natl Acad Sci USA 111:17290–17295. doi:10.1073/pnas.1413204111

Mabb AM, Simon JM, King IF, Lee H-M, An L-K, Philpot BD, Zylka MJ. 2016. Topoisomerase 1 Regulates Gene Expression in Neurons through Cleavage Complex-Dependent and - Independent Mechanisms. PLoS ONE 11:e0156439. doi:10.1371/journal.pone.0156439

Makeyev EV, Zhang J, Carrasco MA, Maniatis T. 2007. The MicroRNA miR-124 Promotes Neuronal Differentiation by Triggering Brain-Specific Alternative Pre-mRNA Splicing. Molecular Cell 27:435–448. doi:10.1016/j.molcel.2007.07.015

McCoy MJ, Fire AZ. 2020. Intron and gene size expansion during nervous system evolution. BMC Genomics 21:360. doi:10.1186/s12864-020-6760-4

McCoy MJ, Paul AJ, Victor MB, Richner M, Gabel HW, Gong H, Yoo AS, Ahn T-H. 2018. LONGO: an R package for interactive gene length dependent analysis for neuronal identity. Bioinformatics 34:i422–i428. doi:10.1093/bioinformatics/bty243

McQuin C, Goodman A, Chernyshev V, Kamentsky L, Cimini BA, Karhohs KW, Doan M, Ding L, Rafelski SM, Thirstrup D, Wiegraebe W, Singh S, Becker T, Caicedo JC, Carpenter AE. 2018. CellProfiler 3.0: Next-generation image processing for biology. PLoS Biol 16:e2005970. doi:10.1371/journal.pbio.2005970

Moore MJ, Zhang C, Gantman EC, Mele A, Darnell JC, Darnell RB. 2014. Mapping Argonaute and conventional RNA-binding protein interactions with RNA at single-nucleotide resolution using HITS-CLIP and CIMS analysis. Nat Protoc 9:263–293. doi:10.1038/nprot.2014.012

Neo WH, Yap K, Lee SH, Looi LS, Khandelia P, Neo SX, Makeyev EV, Su I. 2014. MicroRNA miR-124 Controls the Choice between Neuronal and Astrocyte Differentiation by Fine-tuning Ezh2 Expression. Journal of Biological Chemistry 289:20788–20801. doi:10.1074/jbc.M113.525493

Okano HJ, Darnell RB. 1997. A Hierarchy of Hu RNA Binding Proteins in Developing and Adult Neurons. The Journal of Neuroscience 17:3024–3037. doi:10.1523/JNEUROSCI.17-09-03024.1997

Packer AN, Xing Y, Harper SQ, Jones L, Davidson BL. 2008. The Bifunctional microRNA miR-9/miR-9* Regulates REST and CoREST and Is Downregulated in Huntington’s Disease. Journal of Neuroscience 28:14341–14346. doi:10.1523/JNEUROSCI.2390-08.2008

Partek Inc. 2020. Partek Flow. St. Louis.

Paz I, Kosti I, Ares M, Cline M, Mandel-Gutfreund Y. 2014. RBPmap: a web server for mapping binding sites of RNA-binding proteins. Nucleic Acids Research 42:W361–W367. doi:10.1093/nar/gku406

Plass M, Rasmussen SH, Krogh A. 2017. Highly accessible AU-rich regions in 3’ untranslated regions are hotspots for binding of regulatory factors. PLOS Computational Biology 13:e1005460. doi:10.1371/journal.pcbi.1005460

Rehmsmeier M. 2004. Fast and effective prediction of microRNA/target duplexes. RNA 10:1507–1517. doi:10.1261/rna.5248604

Richner M, Victor MB, Liu Y, Abernathy D, Yoo AS. 2015. MicroRNA-based conversion of human fibroblasts into striatal medium spiny neurons. Nature Protocols 10:1543–1555. doi:10.1038/nprot.2015.102

Robinson MD, McCarthy DJ, Smyth GK. 2010. edgeR: a Bioconductor package for differential expression analysis of digital gene expression data. Bioinformatics 26:139–140. doi:10.1093/bioinformatics/btp616

Scheckel C, Drapeau E, Frias MA, Park CY, Fak J, Zucker-Scharff I, Kou Y, Haroutunian V, Ma’ayan A, Buxbaum JD, Darnell RB. 2016. Regulatory consequences of neuronal ELAV-like protein binding to coding and non-coding RNAs in human brain. eLife 5. doi:10.7554/eLife.10421

Schoenherr C, Anderson D. 1995. The neuron-restrictive silencer factor (NRSF): a coordinate repressor of multiple neuron-specific genes. Science 267:1360–1363. doi:10.1126/science.7871435

Spassov DS, Jurecic R. 2002. Cloning and comparative sequence analysis of PUM1 and PUM2 genes, human members of the Pumilio family of RNA-binding proteinsq 10.

Staahl BT, Tang J, Wu W, Sun A, Gitler AD, Yoo AS, Crabtree GR. 2013. Kinetic Analysis of npBAF to nBAF Switching Reveals Exchange of SS18 with CREST and Integration with Neural Developmental Pathways. Journal of Neuroscience 33:10348–10361. doi:10.1523/JNEUROSCI.1258-13.2013

Sugino K, Hempel CM, Okaty BW, Arnson HA, Kato S, Dani VS, Nelson SB. 2014. Cell-Type-Specific Repression by Methyl-CpG-Binding Protein 2 Is Biased toward Long Genes. Journal of Neuroscience 34:12877–12883. doi:10.1523/JNEUROSCI.2674-14.2014

Sun AX, Yuan Q, Tan S, Xiao Y, Wang D, Khoo ATT, Sani L, Tran H-D, Kim P, Chiew YS, Lee KJ, Yen Y-C, Ng HH, Lim B, Je HS. 2016. Direct Induction and Functional Maturation of Forebrain GABAergic Neurons from Human Pluripotent Stem Cells. Cell Reports 16:1942–1953. doi:10.1016/j.celrep.2016.07.035

Tripathi S, Pohl MO, Zhou Y, Rodriguez-Frandsen A, Wang G, Stein DA, Moulton HM, DeJesus P, Che J, Mulder LCF, Yángüez E, Andenmatten D, Pache L, Manicassamy B, Albrecht RA, Gonzalez MG, Nguyen Q, Brass A, Elledge S, White M, Shapira S, Hacohen N, Karlas A, Meyer TF, Shales M, Gatorano A, Johnson JR, Jang G, Johnson T, Verschueren E, Sanders D, Krogan N, Shaw M, König R, Stertz S, García-Sastre A, Chanda SK. 2015. Meta- and Orthogonal Integration of Influenza “OMICs” Data Defines a Role for UBR4 in Virus Budding. Cell Host & Microbe 18:723–735. doi:10.1016/j.chom.2015.11.002

Truesdell SS, Mortensen RD, Seo M, Schroeder JC, Lee JH, LeTonqueze O, Vasudevan S. 2012. MicroRNA-mediated mRNA Translation Activation in Quiescent Cells and Oocytes Involves Recruitment of a Nuclear microRNP. Scientific Reports 2. doi:10.1038/srep00842

Tsutsui Ken, Tsutsui Kimiko, Sano K, Kikuchi A, Tokunaga A. 2001. Involvement of DNA Topoisomerase IIβ in Neuronal Differentiation. Journal of Biological Chemistry 276:5769–5778. doi:10.1074/jbc.M008517200

Vasudevan S, Steitz JA. 2007. AU-Rich-Element-Mediated Upregulation of Translation by FXR1 and Argonaute 2. Cell 128:1105–1118. doi:10.1016/j.cell.2007.01.038

Vasudevan S, Tong Y, Steitz JA. 2007. Switching from Repression to Activation: MicroRNAs Can Up-Regulate Translation. Science 318:1931–1934. doi:10.1126/science.1149460

Victor MB, Richner M, Hermanstyne TO, Ransdell JL, Sobieski C, Deng P-Y, Klyachko VA, Nerbonne JM, Yoo AS. 2014. Generation of Human Striatal Neurons by MicroRNA-Dependent Direct Conversion of Fibroblasts. Neuron 84:311–323. doi:10.1016/j.neuron.2014.10.016

Victor MB, Richner M, Olsen HE, Lee SW, Monteys AM, Ma C, Huh CJ, Zhang B, Davidson BL, Yang XW, Yoo AS. 2018. Striatal neurons directly converted from Huntington’s disease patient fibroblasts recapitulate age-associated disease phenotypes. Nature Neuroscience. doi:10.1038/s41593-018-0075-7

Visvanathan J, Lee S, Lee B, Lee JW, Lee S-K. 2007. The microRNA miR-124 antagonizes the anti-neural REST/SCP1 pathway during embryonic CNS development. Genes & Development 21:744–749. doi:10.1101/gad.1519107

Vuong JK, Lin C-H, Zhang M, Chen L, Black DL, Zheng S. 2016. PTBP1 and PTBP2 Serve Both Specific and Redundant Functions in Neuronal Pre-mRNA Splicing. Cell Reports 17:2766–2775. doi:10.1016/j.celrep.2016.11.034

Watanabe M, Tsutsui Ken, Tsutsui Kimiko, Inoue Y. 1994. Differential expressions of the topoisomerase IIa and IIb mRNAs in developing rat brain. Neuroscience Research 19:51–57.

Xue Y, Qian H, Hu J, Zhou B, Zhou Y, Hu X, Karakhanyan A, Pang Z, Fu X-D. 2016. Sequential regulatory loops as key gatekeepers for neuronal reprogramming in human cells. Nature Neuroscience 19:807–815. doi:10.1038/nn.4297

Yoo AS, Staahl BT, Chen L, Crabtree GR. 2009. MicroRNA-mediated switching of chromatin-remodelling complexes in neural development. Nature 460:642–646. doi:10.1038/nature08139

Yoo AS, Sun AX, Li L, Shcheglovitov A, Portmann T, Li Y, Lee-Messer C, Dolmetsch RE, Tsien RW, Crabtree GR. 2011. MicroRNA-mediated conversion of human fibroblasts to neurons. Nature 476:228–231. doi:10.1038/nature10323

Yu H, Wang J, Sheng Q, Liu Q, Shyr Y. 2018. beRBP: binding estimation for human RNA-binding proteins. Nucleic Acids Research. doi:10.1093/nar/gky1294

Zhang Y, Liu T, Meyer CA, Eeckhoute J, Johnson DS, Bernstein BE, Nussbaum C, Myers RM, Brown M, Li W, Liu XS. 2008. Model-based Analysis of ChIP-Seq (MACS). Genome Biol 9:R137. doi:10.1186/gb-2008-9-9-r137

Zheng S, Gray EE, Chawla G, Porse BT, O’Dell TJ, Black DL. 2012. PSD-95 is post-transcriptionally repressed during early neural development by PTBP1 and PTBP2. Nature Neuroscience 15:381–388. doi:10.1038/nn.3026

Zhou S, Gao R, Hu W, Qian T, Wang N, Ding G, Ding F, Yu B, Gu X. 2014. miR-9 inhibits Schwann cell migration by targeting Cthrc1 following sciatic nerve injury. Journal of Cell Science 127:967–976. doi:10.1242/jcs.131672

