## Supplemental data for "MiR-124 synergism with ELAVL3 enhances target gene expression to promote neuronal maturity"

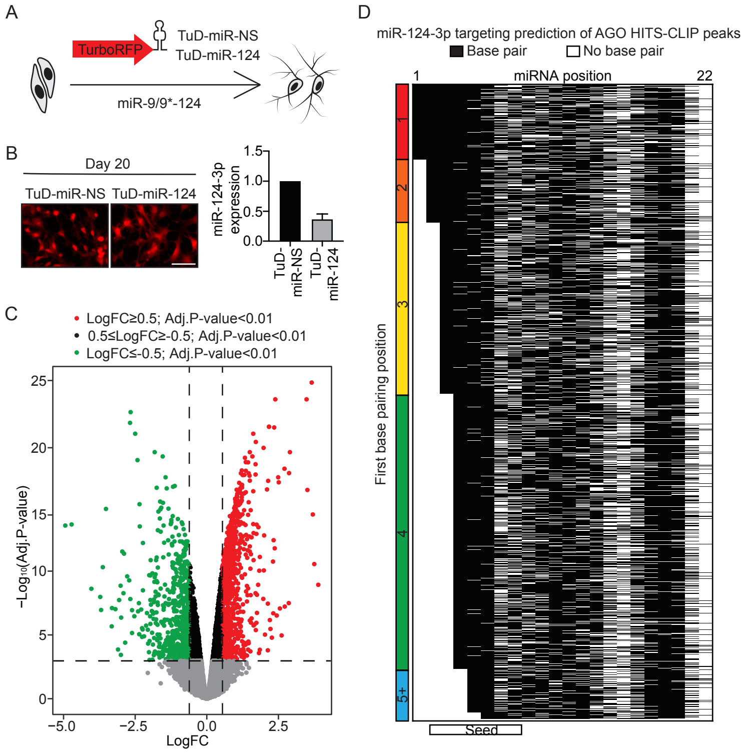


**Figure S1**. MiR-124 target genes during neuronal reprogramming

(A) A schematic diagram of the construct expressing the turbo-RFP reporter attached to the tough decoy (TuD) delivered into miNs.

(B) Left, day 20 miNs showing RFP expression of cells with TuD-miR-NS or TuD-miR-124. Right, Taqman qPCR for mature miR-124-3p showing reduced miR-124 expression upon TuD-miR-124 treatment compared to TuD-miR-NS control. Scale bar = 75 µm.

(C) A volcano plot of DEGs (TuD-miR-124 over the TuD-miR-NS control) in day 20 miNs.

(D) RNAhybrid predictions of base-pairing between miR-123-3p and extracted CLIP sequences of targets from Figure 2 with an energy threshold of -20 kcal/mol or less. The heatmap shows the base-pairing position between miR-124-3p and extracted sequences from AGO HITS-CLIP peaks: base-pairing = black, and gaps or no base-pairing = white.

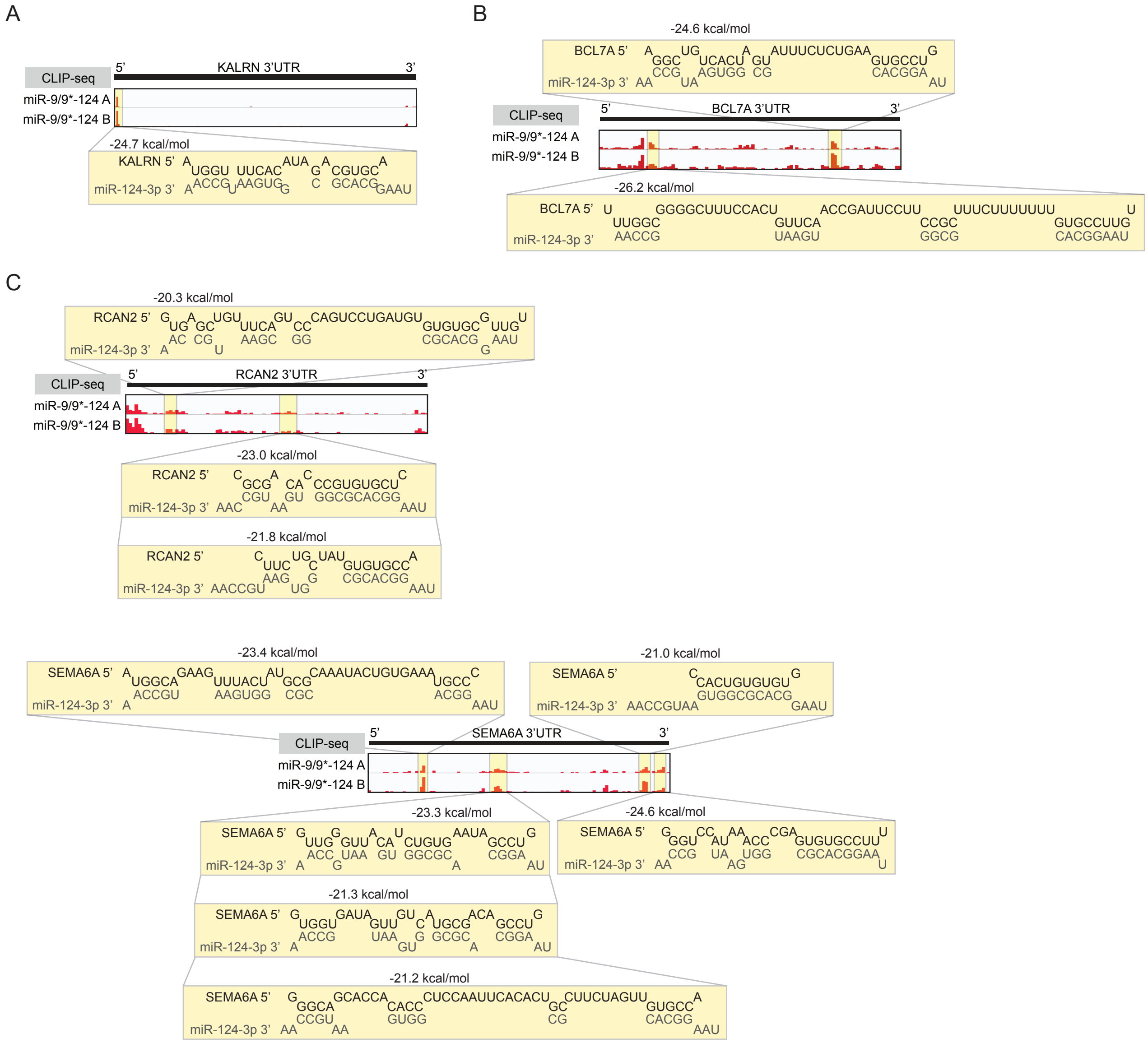


**Figure S2**. Examples of RNAhybrid prediction of miR-124-3p hybridization to enriched 3'UTR HITS-CLIP peak sequences of day 20 upregulated DEGs in miNs

(A) *KALRN* contains one enriched AGO HITS-CLIP peak for miR-124-3p at 3'UTR highlighted in yellow.

(B) *BCL7A* 3'UTR contains two predicted miR-124-3p sites within the AGO HITS-CLIP peaks highlighted in yellow.

(C) *RCAN2* and *SEMA6A* 3'UTRs contain multiple miR-124-3p target sites within the AGO HITS-CLIP peaks highlighted in yellow.

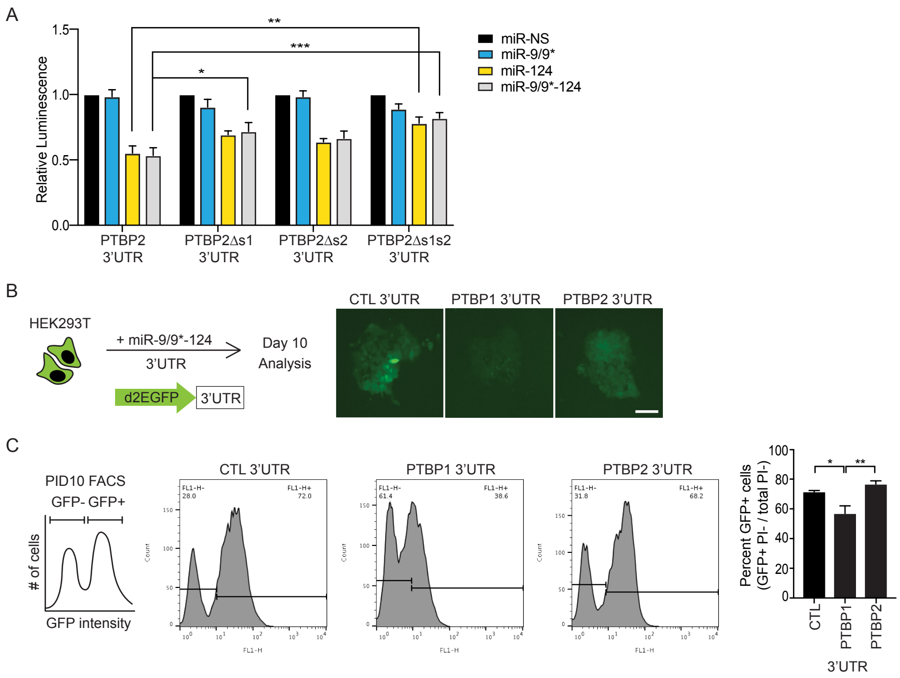


**Figure S3**. MiR-124 target site validation, and assessment of miR-124 promoting PTBP2 expression via targetting PTBP2 3'UTR with the prolonged neurogenic input.

(A) Luciferase assays in HEK293T cells of PTBP2 3'UTR with miR-124 site(s) mutagenized. Luminescence was measured 48 hrs after transfection and normalized to miR-NS control of each condition. Data are represented by mean ± SEM from four independent experiments. Two-way ANOVA followed by Dunnett's test (from left, miR-124 ** P = 0.001; miR-9/9*-124 * P = 0.0104, *** P = 0.0001).

(B) Left, a schematic diagram of the reporter construct and experiments. HEK293T cells were transduced with destabilized EGFP reporter to monitor the 3'UTR activity. Right, representative images of GFP fluorescence of HEK293T cells measured 10 days after miR-9/9*-124 transduction. Scale bar = 100 μm.

(C) Left, schematic of a histogram of flow cytometry analysis with gating of EGFP intensity to separate GFP-negative and GFP-positive cell populations at day 10. Middle, representative images of histograms following flow cytometry. Right, quantification flow cytometry to determine percent GFP-positive, PI-negative cells over the total PI-negative cells. Data represented in mean ± SEM from five independent experiments. One-way ANOVA followed by Tukey's test (from left, * P = 0.0222; ** P = 0.0031).

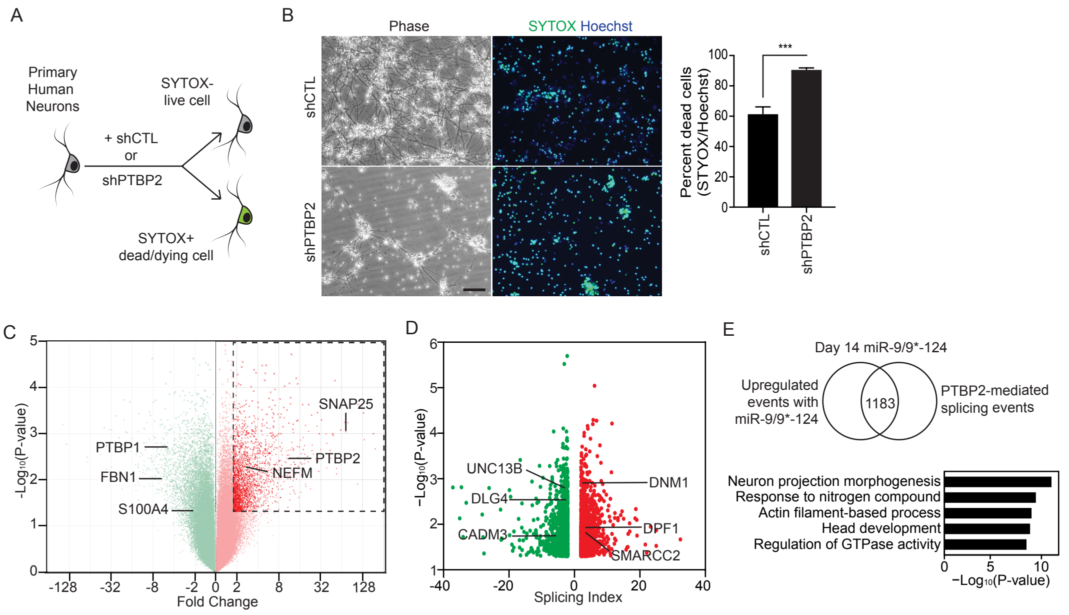


**Figure S4**. PTBP2 plays an essential role during miRNA-mediated neuronal conversion
(A) A schematic diagram of SYTOX assay. Knockdown of PTBP2 with shRNA in primary human neurons resulted in increased cell death measured by SYTOX after 10 days.
(B) Left, representative images of phase contrast and SYTOX staining 10 days after knockdown with CTL or PTBP2 shRNA. Right, quantification of SYTOX-positive cells over total Hoechst-positive cells. Scale bar = 75 μm. Data are represented by mean ± SEM. shGFP n = SYTOX 1237/2073; shPTBP2 n = 1589/1759 scored from independent fields of view. Unpaired t-test (*** P = < 0.0001).
(C) A volcano plot comparing gene expression between miR-9/9*-124 and miR-NS expression on day 14. Red dots within the enclosed box indicate differentially expressed genes (fold change ≥ 1.5, ANOVA $p < 0.05$) that are upregulated in miR-9/9*-124 condition. Examples of non-neuronal and neuronal genes are highlighted in green and red dots, respectively.
(D) A volcano plot of differentially spliced events (splicing index ≥ 2, ANOVA $P < 0.05$) at day 14 comparing shPTBP2 to shCTL in the miR-9/9*-124 expression background. Examples of spliced targets are highlighted with gene names corresponding to the differential splicing events.
(E) Top, target genes of interest were obtained by overlapping upregulated neuronal-associated events in (C) to all differentially spliced events by PTBP2 in (D). Bottom, top biological GO terms associated with the1183 unique events of genes upregulated during neuronal reprogramming and targetted by PTBP2.

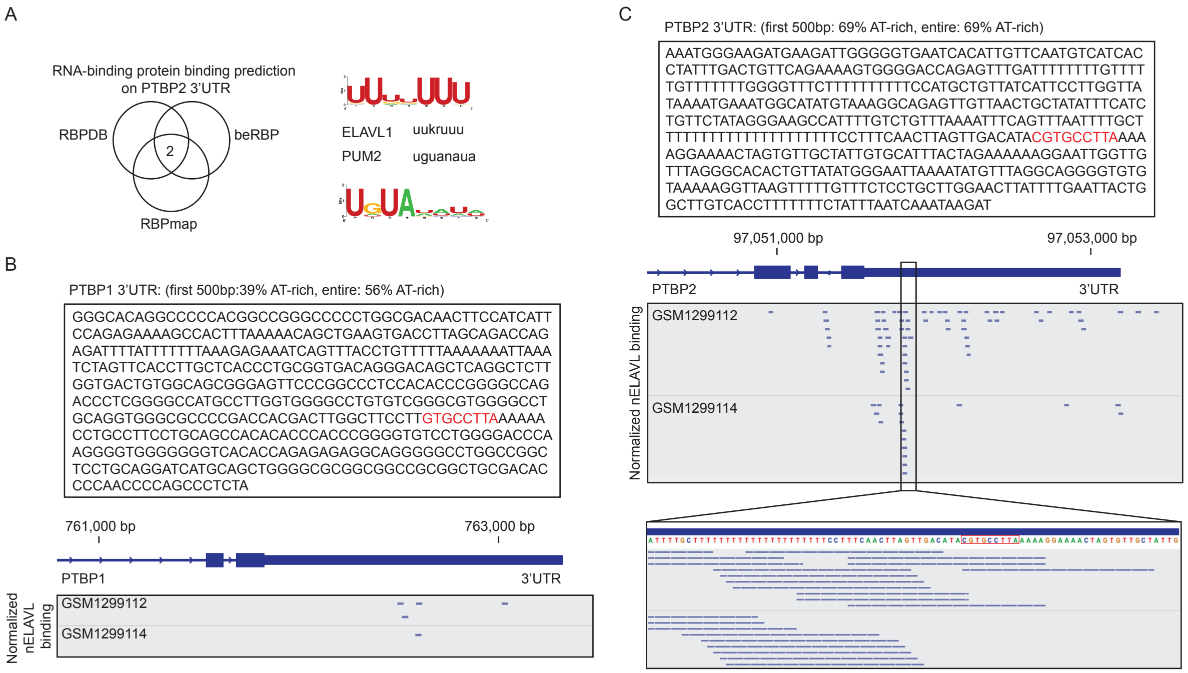


**Figure S5**. nELAVLs bind preferentially to PTBP2 and not PTBP1 3'UTR in the human brain

(A) Left, a Venn diagram overlapping all RNA-binding proteins predicted to bind to PTBP2 3'UTR using three different RNA-binding protein prediction databases. Right, binding motifs on PTBP2 3'UTR of the two predicted candidate RNA-binding proteins (left) based on all predicted motifs on PTBP2 3'UTR from RBPmap.

(B) Top, first 500 bp of PTBP1 3'UTR with the first conserved miR-124 target site in red. Bottom, track views of nELAVL HITS-CLIP of the human brain at PTBP1 3'UTR (GSM129912, GSM129914).

(C) Top, first 500 bp of PTBP2 3'UTR with the first conserved miR-124 target site in red. Bottom, track views of nELAVL HITS-CLIP of the human brain at PTBP2 3'UTR (GSM129912, GSM129914). The enclosed red box denotes the miR-124 target site.

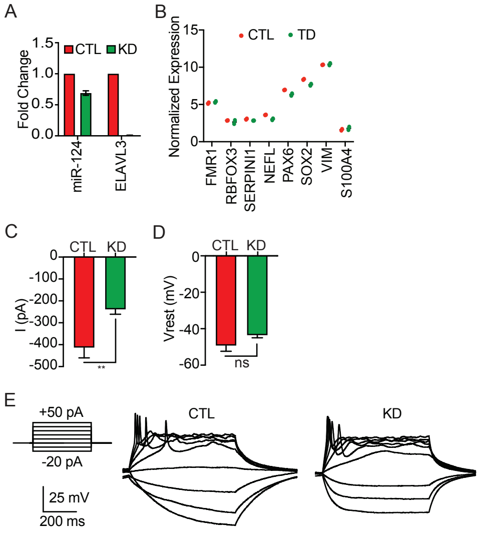


**Figure S6**. Neuronal long genes are targets of miR-124 and nELAVLs

(A) RT-qPCR for mature miR-124 expression and ELAVL3, showing reduced miR-124 and ELAVL3 expression upon KD treatment compared to CTL.

(B) Expression of genes associated with neuronal and neuronal progenitor identity in CTL and KD HNs.

(C) Peak inward sodium current amplitude of CTL and KD HNs. Data are represented in ± SEM from seven recorded cells from each condition. Two-tailed unpaired t-test (** P = 0.005).

(D) Resting membrane potential of CTL and KD HNs. Data are represented in ± SEM from seven recorded cells from each condition. Two-tailed unpaired t-test (ns P = 0.1191).

(E) Representative current-clamp traces of CTL and KD HNs.

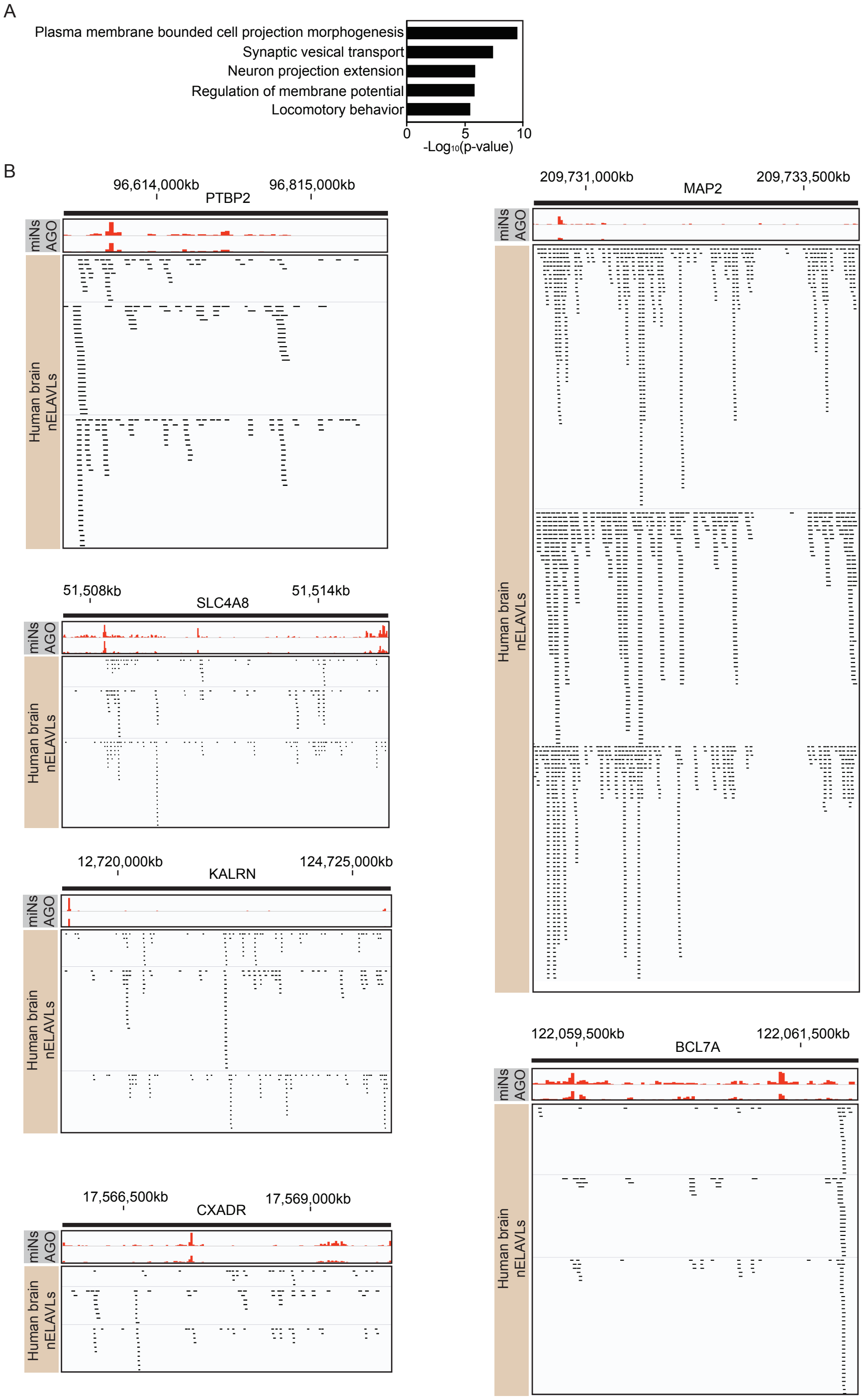


**Figure S7.** Identified neuronal transcripts that are bound by AGO and nELAVLs in HNs
(A) Top biological GO terms associated with the 77 genes that are predicted to be targeted by both AGO-miR-124 and ELAVL3, based on AGO HITS-CLIP in miNs, nELAVL HITS-CLIP in human brain, and HN RNA-seq with miR-124 and ELAVL3 knockdown.
(B) Track views of AGO HITS-CLIP in miNs and nELAVL HITS-CLIP in the human brain (GSM1299112, GSM1885137, GSM129911) of a select neuronal transcript 3'UTRs identified to be targets of AGO-miR-124 in miNs and HNs with nELAVL binding.

| Gene name | logFC | adj.P.Val |
|---|---|---|
| ACAD10 | 2.69E-01 | 1.25E-03 |
| ACER2 | -6.41E-01 | 2.75E-06 |
| ALAS1 | 4.42E-01 | 6.48E-08 |
| ANGPTL1 | 2.62E-01 | 1.11E-04 |
| ANK3 | -1.06E+00 | 4.61E-12 |
| APBA1 | -3.58E-01 | 5.20E-03 |
| APOE | 3.11E-01 | 2.59E-04 |
| ARHGAP20 | -2.58E-01 | 1.14E-03 |
| ARHGEF7 | -2.45E-01 | 3.73E-04 |
| ATP6V0A1 | -3.55E-01 | 4.02E-06 |
| ATP6V1G2 | -7.71E-01 | 1.91E-11 |
| BCL7A | -5.14E-01 | 2.80E-04 |
| BDH2 | 2.56E-01 | 1.23E-04 |
| BMF | -9.39E-01 | 6.06E-10 |
| C14orf132 | 2.55E-01 | 8.72E-05 |
| CAMK4 | -9.30E-01 | 6.86E-10 |
| CAMSAP1 | -4.72E-01 | 1.66E-04 |
| CDK15 | -8.77E-01 | 3.34E-08 |
| CDS1 | -1.53E+00 | 4.75E-08 |
| CDS2 | -1.55E-01 | 6.90E-03 |
| CEP85L | -5.40E-01 | 4.77E-09 |
| CERS6 | -3.93E-01 | 1.22E-04 |
| CFI | 1.73E-01 | 5.31E-03 |
| CHRDL1 | -3.77E-01 | 4.16E-03 |
| CLCN4 | -2.56E-01 | 1.76E-03 |
| CLCN6 | -6.20E-01 | 5.98E-10 |
| CLDN11 | 4.59E-01 | 1.06E-07 |
| CLUL1 | -1.53E+00 | 4.41E-16 |
| CNNM1 | -8.45E-01 | 1.74E-05 |
| CNTN1 | -1.15E+00 | 2.06E-05 |
| COX6C | 4.76E-01 | 1.44E-08 |
| CPE | 2.48E-01 | 1.51E-03 |
| CRY1 | -8.57E-01 | 1.47E-09 |
| CRY2 | -6.05E-01 | 1.25E-07 |
| CXADR | -1.06E+00 | 1.24E-10 |
| DCLK1 | -8.70E-01 | 4.73E-13 |
| DHX30 | -3.00E-01 | 8.47E-05 |
| DKK2 | -5.74E-01 | 1.69E-06 |
| DNAJB5 | -6.00E-01 | 2.77E-09 |
| DOCK3 | -4.80E-01 | 2.08E-04 |
| DTNA | -3.15E-01 | 4.16E-05 |
| DUSP26 | -1.36E+00 | 3.71E-05 |
| DUSP8 | -1.98E+00 | 3.49E-07 |
| EEF1A2 | -1.95E+00 | 4.25E-08 |
| EFR3B | -8.75E-01 | 6.22E-11 |
| ENOX1 | -7.19E-01 | 1.34E-03 |
| EPB41 | -3.94E-01 | 2.63E-03 |
| EPHA4 | 2.73E-01 | 9.07E-04 |
| FADS2 | 2.97E-01 | 2.50E-04 |
| FAIM2 | 2.85E-01 | 8.55E-04 |
| FAM110B | -2.19E-01 | 1.48E-03 |
| FAM13C | -3.07E-01 | 4.61E-06 |
| FAM184A | -8.65E-01 | 3.96E-06 |
| FILIP1 | -7.07E-01 | 3.53E-09 |
| FKBP5 | -3.28E-01 | 2.16E-03 |
| FRMD4A | -4.16E-01 | 3.84E-05 |
| FSD1L | -2.44E-01 | 9.50E-03 |
| GABRE | -2.10E-01 | 1.21E-03 |
| GAD1 | -9.23E-01 | 4.01E-07 |
| GDF11 | -1.93E-01 | 9.47E-03 |
| GDF5 | -1.90E+00 | 1.49E-10 |
| GPM6A | -8.05E-01 | 9.82E-03 |
| GPNMB | 1.55E-01 | 7.24E-03 |
| GRIA2 | -9.12E-01 | 1.05E-03 |
| GRIA4 | -1.44E+00 | 1.39E-04 |

| Gene name | logFC | adj.P.Val |
|---|---|---|
| MDM1 | -2.89E-01 | 1.26E-04 |
| MFHAS1 | -3.46E-01 | 4.90E-03 |
| MGAT4C | -9.34E-01 | 1.15E-03 |
| MITF | -3.52E-01 | 5.53E-03 |
| MRPL35 | 2.00E-01 | 2.01E-03 |
| MTHFR | -4.84E-01 | 2.25E-07 |
| NEFM | -7.36E-01 | 3.27E-06 |
| NINJ1 | 3.22E-01 | 1.49E-05 |
| NLGN2 | 2.99E-01 | 8.63E-03 |
| OGFRL1 | -3.40E-01 | 7.90E-04 |
| OSBPL1A | 4.04E-01 | 2.55E-06 |
| OXR1 | -2.10E-01 | 5.70E-04 |
| PASK | -7.59E-01 | 2.82E-03 |
| PBX3 | -9.96E-01 | 1.24E-09 |
| PCDH7 | 4.33E-01 | 8.56E-06 |
| PCLO | -1.42E+00 | 1.78E-04 |
| PELP1 | -4.45E-01 | 2.19E-03 |
| PER3 | -5.22E-01 | 8.66E-06 |
| PFKM | 4.86E-01 | 9.38E-10 |
| PGM2L1 | 3.27E-01 | 2.03E-05 |
| PHKA1 | -4.32E-01 | 4.44E-04 |
| PHLPP1 | -3.60E-01 | 1.38E-03 |
| PHYHIPL | -7.93E-01 | 1.30E-03 |
| PLD6 | -6.71E-01 | 1.74E-03 |
| PNISR | -2.62E-01 | 1.39E-05 |
| PPA2 | 2.47E-01 | 1.13E-04 |
| PPM1D | -3.89E-01 | 1.96E-04 |
| PPM1H | -9.24E-01 | 3.05E-11 |
| PPP1R9A | -1.14E+00 | 2.39E-11 |
| PTBP2 | 2.45E-01 | 2.20E-04 |
| PTN | -8.37E-01 | 8.82E-11 |
| RALGAPA1 | -2.61E-01 | 4.19E-03 |
| RALGPS1 | -5.08E-01 | 1.06E-03 |
| RANBP3L | -9.82E-01 | 8.23E-12 |
| RAPGEF4 | -1.70E+00 | 6.51E-09 |
| RASL10B | -6.35E-01 | 8.25E-04 |
| RCAN2 | -7.31E-01 | 1.90E-12 |
| RIC3 | -1.50E+00 | 6.66E-08 |
| RIMBP2 | -1.19E+00 | 9.69E-05 |
| RIMS2 | -3.26E-01 | 2.58E-03 |
| RNF157 | -4.66E-01 | 6.72E-06 |
| RNFT2 | -8.03E-01 | 3.39E-11 |
| RRAGD | -1.64E+00 | 7.49E-15 |
| SAMD12 | -1.21E+00 | 3.44E-09 |
| SCARB1 | 4.19E-01 | 6.87E-03 |
| SCN1A | -7.43E-01 | 8.57E-08 |
| SCN2A | -2.24E-01 | 4.95E-03 |
| SCN9A | -4.44E-01 | 4.53E-04 |
| SDHA | 4.27E-01 | 1.05E-08 |
| SEMA6A | -1.15E+00 | 1.18E-09 |
| SENP7 | -2.45E-01 | 3.82E-03 |
| SESN3 | -3.27E-01 | 4.58E-04 |
| SFMBT2 | -9.75E-01 | 2.02E-03 |
| SFRP4 | 4.41E-01 | 4.72E-06 |
| SH3BP2 | 3.52E-01 | 2.16E-04 |
| SLC12A7 | -6.84E-01 | 7.60E-04 |
| SLC25A11 | 2.73E-01 | 4.66E-05 |
| SLC25A13 | 4.31E-01 | 1.64E-07 |
| SLC2A8 | -8.54E-01 | 2.39E-03 |
| SLC4A8 | 3.30E-01 | 2.34E-04 |
| SLC7A14 | -3.73E-01 | 3.33E-05 |
| SLITRK4 | -6.47E-01 | 2.55E-03 |
| SMARCB1 | -4.41E-01 | 1.38E-04 |
| SNAP25 | -5.52E-01 | 5.29E-10 |
| SPARCL1 | -7.89E-01 | 1.32E-04 |

| | | |
|---|---|---|
| HMGCS1 | -5.84E-01 | 3.64E-11 |
| HTR2B | -6.42E-01 | 6.61E-03 |
| IGF1 | 2.00E-01 | 1.21E-03 |
| IGSF10 | -4.78E-01 | 1.99E-04 |
| INPP1 | 3.61E-01 | 7.75E-06 |
| IRF6 | -2.34E+00 | 3.13E-14 |
| ITM2A | -5.17E-01 | 2.47E-03 |
| KALRN | -2.41E-01 | 3.41E-03 |
| KAT2B | -3.56E-01 | 3.43E-03 |
| KCNK1 | -6.44E-01 | 9.90E-03 |
| KIAA1109 | -3.48E-01 | 3.91E-05 |
| KIAA1217 | -4.34E-01 | 3.00E-06 |
| KIF5A | -4.88E-01 | 3.85E-09 |
| KIF5C | -9.14E-01 | 1.40E-11 |
| KLLN | -4.85E-01 | 7.94E-04 |
| KPNA5 | -4.18E-01 | 8.78E-06 |
| LCOR | -3.31E-01 | 3.87E-05 |
| LDB2 | -2.28E-01 | 5.54E-03 |
| LDHB | 4.57E-01 | 4.91E-09 |
| LMBR1 | 3.07E-01 | 1.54E-03 |
| LRRN3 | -1.62E+00 | 1.43E-15 |
| MAGI2 | -7.78E-01 | 1.74E-09 |
| MAMDC2 | 3.69E-01 | 5.44E-04 |
| MAN1C1 | -3.73E-01 | 2.77E-06 |
| MANSC1 | -3.27E-01 | 2.30E-03 |
| MAP2 | -2.94E-01 | 6.73E-05 |
| MAP3K14 | -3.27E-01 | 4.81E-03 |
| MAPRE3 | -5.43E-01 | 3.36E-04 |
| MBP | -1.18E+00 | 5.45E-11 |
| MCM3 | -1.75E-01 | 9.94E-03 |
| MCOLN1 | -3.55E-01 | 1.23E-03 |

| | | |
|---|---|---|
| SPP1 | -8.26E-01 | 3.11E-07 |
| STAC2 | 4.69E-01 | 2.91E-03 |
| STMN2 | 2.14E-01 | 3.39E-03 |
| STRBP | -6.99E-01 | 4.99E-07 |
| STX1B | -2.83E-01 | 6.24E-03 |
| SULT1C4 | -5.37E-01 | 1.42E-05 |
| SV2A | 1.74E-01 | 4.14E-03 |
| SYBU | -7.80E-01 | 3.87E-09 |
| TBC1D9 | -3.30E-01 | 5.43E-03 |
| THSD7A | -6.27E-01 | 2.14E-03 |
| TKT | 2.56E-01 | 4.20E-05 |
| TLE1 | -4.92E-01 | 1.57E-05 |
| TM7SF3 | 2.20E-01 | 5.34E-03 |
| TMEM63B | 4.40E-01 | 9.37E-04 |
| TMEM8B | -3.26E-01 | 5.29E-03 |
| TMOD2 | -4.14E-01 | 2.88E-06 |
| TNFRSF14 | -4.38E-01 | 4.31E-04 |
| TNFRSF21 | 4.14E-01 | 3.92E-06 |
| TRAPPC9 | -3.39E-01 | 6.05E-03 |
| TRPC3 | -1.97E+00 | 1.84E-08 |
| TSHZ1 | -7.13E-01 | 5.06E-09 |
| TSPAN14 | 2.02E-01 | 3.98E-03 |
| TSPAN7 | -1.55E+00 | 4.65E-04 |
| TXLNB | -1.35E+00 | 3.82E-13 |
| UPF3B | 2.19E-01 | 8.66E-03 |
| UQCRC1 | 3.26E-01 | 1.40E-06 |
| WDR7 | -3.75E-01 | 1.51E-05 |
| ZNF518A | -3.87E-01 | 2.06E-04 |
| ZNF577 | -3.15E-01 | 8.04E-04 |
| ZNF738 | -2.97E-01 | 2.23E-03 |
| ZSWIM7 | -3.30E-01 | 1.35E-04 |

Table S1. MiR-124-responsive genes that harbor AGO-enriched peaks in miNs
List of upregulated genes during conversion harboring AGO-enriched peaks and are not upregulated upon miR-124 knockdown in miNs(TuD-miR-124 /TuD-miR-NS; $Log_2FC \geq -0.5$; adj. P.value > 0.01).

| | | | | | | |
|---|---|---|---|---|---|---|
| SEPT4 | ANK1 | ATP9A | CALR | CNTN1 | DIP2B | FAIM2 |
| AAK1 | ANK2 | ATPAF1 | CAMK2D | COA5 | DIS3L2 | FAM102A |
| AASDHPPT | ANKRD20A11P | ATXN1 | CAMK2N1 | COL1A2 | DJA2 | FAM122C |
| AASS | ANKRD36C | ATXN10 | CAMKK1 | COL21A1 | DJB6 | FAM126B |
| ABCA1 | ANKRD44 | AVL9 | CAMSAP1 | COL24A1 | DJC5 | FAM135A |
| ABCA12 | ANKRD46 | AXIN2 | CAMTA1 | COL3A1 | DJC6 | FAM13B |
| ABCA7 | ANO8 | B3GAT3 | CAPZB | COL4A5 | DJC7 | FAM13C |
| ABCB4 | ANP32A | B3GLCT | CBX5 | COL5A2 | DKK3 | FAM169A |
| ABCC6P1 | ANXA11 | B4GALT3 | CCBE1 | COLEC12 | DLD | FAM196B |
| ABCC9 | ANXA6 | BAALC | CCDC136 | COLGALT2 | DLEU2 | FAM198B |
| ABCD2 | AP1B1 | BAG6 | CCDC149 | COMMD2 | DLG4 | FAM20C |
| ABCD3 | AP1M1 | BAZ2B | CCDC64 | COPG2 | DLG5 | FAM234B |
| ABCG4 | AP1S2 | BBS9 | CCDC7 | COX6C | DMPK | FAM49B |
| ABHD6 | AP2S1 | BCAS1 | CCDC74A | CPEB3 | DMXL2 | FAM65B |
| ABLIM3 | AP5M1 | BCL7A | CD109 | CPEB4 | DNM1 | FAM66E |
| ABR | APBB1 | BCL9 | CD14 | CPQ | DNM3 | FAM69B |
| ABTB1 | APC | BDH2 | CD36 | CPSF6 | DNPH1 | FAM96A |
| AC083843.4 | APCDD1 | BEND5 | CD74 | CREB1 | DOCK10 | FAP |
| ACAD11 | APLP2 | BEND6 | CDH18 | CREM | DOCK4 | FAR2 |
| ACAD8 | APOL4 | BLID | CDH6 | CRLS1 | DOK5 | FAR2P2 |
| ACADM | APOO | BMP1 | CDK14 | CRMP1 | DOT1L | FASTKD1 |
| ACAN | APP | bP-21264C1.1 | CDK15 | CROT | DPF1 | FAT4 |
| ACAP3 | AQP9 | BRD3 | CDK16 | CS | DPP3 | FBLN2 |
| ACAT1 | ARFGEF1 | BRE | CDK18 | CSRNP3 | DPT | FBXL17 |
| ACBD5 | ARG2 | BRI3BP | CDKL1 | CTB-52I2.4 | DPYSL2 | FBXO9 |
| ACER2 | ARHGAP12 | BRMS1L | CDS1 | CTPS2 | DPYSL4 | FCMR |
| ACER3 | ARHGAP20 | BRWD3 | CEACAM1 | CTSO | DROSHA | FER |
| ACSF2 | ARHGAP26 | BTRC | CEMIP | CTSS | DST | FGD4 |
| ACSL4 | ARHGAP32 | C10orf10 | CENPW | CUL3 | DT | FGD6 |
| ACTR10 | ARHGAP42 | C11orf49 | CEP85L | CXADR | DUSP23 | FGF13 |
| ACTR3B | ARHGEF10L | C12orf75 | CFI | CXCL12 | DUSP6 | FH |
| ACVR1B | ARL5A | C14orf37 | CH17-472G23.4 | CYB5R4 | dydorbo | FHOD3 |
| ACVR2A | ARMC9 | C15orf39 | CHD4 | CYFIP2 | DYNC1I2 | FIBIN |
| ACVR2B | ARRDC4 | C15orf57 | CHFR | CYGB | DYRK2 | FILIP1 |
| ADAM17 | ARSF | C16orf45 | CHI3L2 | CYP3A4 | EDIL3 | FKBP2 |
| ADAM19 | ARV1 | C18orf54 | CHR3 | CYP3A5 | EDNRA | flawfybo |
| ADAM22 | ASAH1 | C1orf220 | CHRM2 | CYP3A7 | EEF2K | FN3K |
| ADAM23 | ASAP1 | C1QTNF1 | CHST10 | CYP46A1 | EEPD1 | FNDC1 |
| ADAMTS12 | ASB9 | C1QTNF6 | CHST11 | CYTH2 | EFR3B | FNDC4 |
| ADARB1 | ATAD2B | C2orf27A | CIITA | DAAM2 | EHBP1 | FNDC5 |
| ADCY1 | ATAT1 | C4A | CISH | DAB1 | EIF5A2 | FOXN3 |
| ADD2 | ATF6B | C4B | CIT | DAPK1 | ELN | FRAS1 |
| ADGRA3 | ATF7IP | C4orf27 | CKB | daseebo | EMCN | FRK |
| ADGRL1 | ATG4D | C8orf34 | CLASP2 | DCBLD1 | EML6 | FRMD4A |
| ADGRL2 | ATP13A2 | C9orf3 | CLCN4 | DCLK1 | EMP2 | FRMD5 |
| ADK | ATP1A3 | CABLES1 | CLEC16A | DCLK2 | EN1 | FAIM2 |
| AES | ATP1B1 | CABLES2 | CLNS1A | DCTPP1 | ENPP5 | FAM102A |
| AF131217.1 | ATP2B4 | CABP2 | CLSTN3 | DCUN1D4 | ENSA | FAM122C |
| AF186192.5 | ATP5A1 | CAC1A | CLTA | DDAH1 | EPB41L1 | FAM126B |
| AGAP1 | ATP5B | CAC1G | CLTC | DDIT4L | EPB41L3 | FAM135A |
| AGAP2 | ATP5C1 | CAC2D3 | CLU | DDR1 | EPHA4 | FAM13B |
| AIFM1 | ATP5G2 | CACHD1 | CLUH | DDT | EPHB2 | FAM13C |
| AIG1 | ATP5L | CACNB2 | CLUL1 | DDX39B | EPHX2 | FAM169A |
| AK5 | ATP6AP1 | CACNB4 | CLYBL | DGCR8 | EPS15 | FAM196B |
| ALAD2 | ATP6V0A1 | CACNG7 | CNIH2 | DHODH | ETS1 | FAM198B |
| ALG9 | ATP6V0E2 | CADM1 | CNIH3 | DHX15 | EVL | FAM20C |
| AMPD3 | ATP6V1B2 | CADM3 | CNKSR2 | DHX30 | EXOC1 | FAM234B |
| AMPH | ATP6V1C1 | CALM1 | CNOT8 | DHX9 | EXOC4 | FAM49B |
| ANGEL2 | ATP6V1G2-DDX39B | CALM3 | CNTFR | DICER1 | FABP3 | FAM65B |

| | | | | | | |
|---|---|---|---|---|---|---|
| FRMD5 | HLF | KIF26B | MAP2 | MYLK | NEFM | PCBP3 |
| FRMPD3 | HMG20A | KIF3B | MAP2K6 | MYO1B | NEGR1 | PCBP4 |
| FRY | HMGB1 | KIF3C | MAP3K12 | MYO5A | NEK10 | PCCA |
| FYB | HNRNPA3P3 | KIF5C | MAP3K13 | MYOZ2 | NELFCD | PCCB |
| GAB1 | HOMER1 | KLC1 | MAP4K3 | NBEA | NET1 | PCDHA9 |
| GABRA3 | HOXA10 | KLF3 | MAPK10 | NBEAP1 | NETO2 | PCDHB16 |
| GABRE | HOXA13 | KLHL13 | MAPRE2 | NCALD | NEURL4 | PCDHB18P |
| GABRR1 | HR | KLRC4-KLRK1 | MAPRE3 | NCAM1 | NFAT5 | PCDHB8 |
| GAD1 | HRAT17 | KMT5B | MAPT | NCOA1 | NFYC | PCDHGC3 |
| GALC | HSBP1 | KP5 | MATR3 | NDN | NID2 | PCMTD1 |
| GALNT11 | HSD17B4 | LACC1 | mawgar | NDRG1 | NIPSP3A | PCSK5 |
| GALNT14 | HSP90B1 | LACTB | MBOAT1 | NDRG2 | NKIRAS1 | PCSK7 |
| GALNT5 | HSPA12A | LAMB1 | MBTD1 | NDRG4 | NLGN4X | PCYOX1L |
| GAS1RR | HSPB7 | LARGE | MCC | NDUFAF5 | NLK | PDCL |
| GAS7 | HTT | LBH | MCM4 | NDUFS1 | NNT | PDE1A |
| GBP2 | HUNK | LCN | MCOLN3 | NDUFS2 | NOL4L | PDE3A |
| GDF11 | INA | LEPR | MCTP1 | NDUFV2 | NOMO3 | PDE4B |
| GGCT | IAH1 | LGALSL | MDH1B | NECAB3 | NONO | PDGFD |
| GIPC1 | IBTK | LIMCH1 | ME3 | NEFL | NPNT | PDGFRA |
| GLI3 | IDH2 | LINC00673 | MEG3 | NEFM | NRCAM | PDK1 |
| GLUL | IGFBP2 | LINC00702 | MEGF8 | NEGR1 | NREP | PDP1 |
| GNB5 | IGFBP3 | LINC00886 | MEIS3 | NEK10 | NRG1 | PDS5B |
| GNG2 | IGFL4 | LINC01091 | MEMO1 | NELFCD | NRP1 | PDZRN3 |
| GP1BB | IGK | LINC01535 | MEST | NET1 | NSF | PELI2 |
| GPAA1 | IGSF10 | LINC01621 | MET | NETO2 | NSFP1 | PFKM |
| GPCPD1 | IKZF2 | LLGL1 | METTL9 | NEURL4 | NSG1 | PFN2 |
| GPI | IL17RB | LMBR1 | MFAP2 | NFAT5 | NSL1 | PGD |
| GPM6B | IL17RD | LMBR1L | MGAT5 | NFYC | NSMF | PGM2L1 |
| GPR155 | IL6R | LMO7 | MGP | NID2 | NT5C2 | PHACTR1 |
| GPR162 | IL7 | LOC100507002 | MGST3 | NIPSP3A | NT5C3A | PHACTR3 |
| GPR173 | IMMT | LOC101927686 | MICAL2 | NKIRAS1 | NT5DC1 | PHKA1 |
| GPR75-ASB3 | INPP4B | LOC101929710 | MICU1 | NLGN4X | NTN4 | PHYHIPL |
| GPRC5B | INPP5A | LOC102723373 | MICU3 | NLK | NTNG1 | PI4KA |
| GRAMD1B | INSRR | LOC102723769 | MID1 | NNT | NTRK1 | PI4KAP1 |
| GRAMD4 | IQCJ | LOC105378663 | MINOS1P1 | NOL4L | NTRK3 | PI4KAP2 |
| GREM1 | ITGA11 | LOC220729 | MIR22HG | NOMO3 | NUAK1 | PIAS4 |
| GS | ITGA2 | LOC645513 | MIR29A | NONO | NUDCD3 | PIK3CB |
| GSR | ITGA5 | LOC728730 | MIR4500HG | NPNT | NUDT22 | PIK3R1 |
| GSTM2 | ITGAV | LPAR6 | MIR573 | NRCAM | NUDT3 | PIK3R2 |
| GTF2I | ITM2A | LPCAT1 | MIR99AHG | NREP | OBSCN | PIK3R3 |
| GTF2IP1 | ITSN1 | LPP-AS2 | MLIP | MYLK | OGDH | PIN1 |
| GTF3C1 | JARID2 | LRP4 | MLLT11 | MYO1B | OGFRL1 | PIP4K2A |
| GTF3C2 | JPH2 | LRP5 | MMD | MYO5A | OIP5-AS1 | PIP5K1C |
| GUCY1B3 | KALRN | LRRC32 | MMP16 | MYOZ2 | ORMDL3 | PITPNM1 |
| GULP1 | KCB2 | LRRC40 | MOB4 | NBEA | OSBP2 | PKIA |
| GXYLT2 | KCNC4 | LRRC49 | moymoyby | NBEAP1 | OSBPL1A | PKIG |
| H2AFY | KCNH5 | LRRK2 | MPC1 | NCALD | OSCP1 | PKM |
| H2AFY2 | KCNK2 | LSAMP | MREG | NCAM1 | OSR1 | PLA2G15 |
| H3F3B | KCNMA1 | LUCAT1 | MROH7 | NCOA1 | OXTR | PLA2G4A |
| HABP4 | KDM4C | LYNX1 | MRPL35 | NDN | P1L5 | PLAGL1 |
| HAGH | KDM5B | LYST | MRPS7 | NDRG1 | P4HTM | PLCB1 |
| HDAC5 | KHDRBS1 | maby | MSI1 | NDRG2 | PAFAH1B2 | PLCB4 |
| HDAC9 | KIAA0391 | MAFB | MT1F | NDRG4 | PAK1 | PLCD4 |
| HECW1 | KIAA0895L | MAGI1 | MTMR1 | NDUFAF5 | PANK1 | PLCH1 |
| HEPH | KIAA0922 | MAGI3 | MTMR9 | NDUFS1 | PARM1 | PLD1 |
| HIST4H4 | KIAA1109 | MAMDC2 | MTSS1L | NDUFS2 | PARP8 | PLEKHA2 |
| HK1 | KIAA1644 | MAN1C1 | MTUS1 | NDUFV2 | PB | PLEKHA5 |
| HK2 | KIF1B | MAOA | MXRA5 | NECAB3 | PBX1 | PLIN2 |
| HLA-DRA | KIF21A | MAP1B | MYH10 | NEFL | PBX3 | PLK2 |
| PLPPR4 | RALGAPA1 | SCP2 | SLC9B2 | SVILP1 | TP53I11 | WARS2-IT1 |
| PLS3 | RALGAPA2 | SCRG1 | SLIT2 | SYNE1 | TP53I3 | WASF3 |
| pluspar | RALGDS | SDCCAG8 | SMAD1 | SYNGR3 | TP63 | WBP2 |

| | | | | | | |
|---|---|---|---|---|---|---|
| PLXDC1 | RALGPS1 | SDHA | SMARCA2 | SYNJ1 | TPCN1 | WDR37 |
| PLXDC2 | RANBP3L | SDHAP3 | SMARCA4 | SYNJ2 | TPD52L1 | WDR41 |
| PLXNC1 | RAP1GAP2 | SDK2 | SMARCC2 | SYNPO2 | TRAK1 | WDR63 |
| PLXND1 | RASA3 | SEC14L1 | SMG6 | SYPL2 | TRAPPC2 | WDR7 |
| PMAIP1 | RASGRP3 | SEC61A2 | SMIM10L2A | SYT1 | TRDMT1 | weewerby |
| POLR2E | RASL10B | SECTM1 | SNN | T14 | TRERF1 | WHSC1 |
| POLR3A | RBM12B | SEMA3D | SNRPA | TADA2A | TRIB1 | WWP2 |
| POLR3F | RBM24 | SEMA6B | SNRPA1 | TANC1 | TRIM16L | XPNPEP1 |
| POMGNT1 | RBMX | SEMA6D | SOCS2 | TAPT1 | TRIM2 | YAF2 |
| POMK | RERE | SENP7 | SOCS2-AS1 | TBC1D12 | TRIM24 | yara |
| POR | REREP3 | SEPN1 | SOGA3 | TBC1D8 | TRMT2B | YLPM1 |
| POSTN | REV3L | SEPT7P2 | SORBS1 | TBCB | TRPC3 | YPEL3 |
| PPARD | RFTN1 | SEPW1 | SORL1 | TBCK | TSPAN13 | YWHAE |
| PPARGC1A | RGL1 | SERPINF1 | SOX4 | TCIRG1 | TSPAN15 | ZC2HC1A |
| PPFIA4 | RGS16 | SERPINI1 | SPAG17 | TCN2 | TSPAN18 | ZC3H12B |
| PPIA | RGS3 | SETBP1 | SPARC | TCONS_l2_00002184 | TSPAN3 | ZCCHC2 |
| PPIAL4C | RGS4 | SFRP4 | SPARCL1 | TCONS_l2_00026703 | TSPAN5 | ZDHHC13 |
| PPP1R12B | RGS8 | SGIP1 | SPATA13 | TCONS_l2_00030954 | TSPAN7 | ZDHHC15 |
| PPP1R14B | RHOBTB1 | SH3BP2 | SPATA20 | TDO2 | TSSC1 | ZDHHC16 |
| PPP1R21 | RIMS2 | SH3BP5 | SPATA6 | TENM4 | TTC13 | ZDHHC9 |
| PPP2R1B | RIT1 | SH3GLB2 | SPC3 | TFAM | TTC19 | ZEB1-AS1 |
| PPP2R2B | RMDN2 | SHOX2 | SPIDR | TGFBI | TTC33 | ZFAND5 |
| PPP2R2D | RNF150 | SHTN1 | SPOCK1 | TGM2 | TTLL6 | ZFAND6 |
| PPP2R3A | RNF38 | SIAE | SPRY1 | THRA | TTYH3 | ZFHX4 |
| PPP5C | RNFT2 | SIPA1L3 | SPRYD3 | THRB | TUBA4A | ZFP14 |
| PQLC2L | ROR1 | SIRPA | SPSB1 | THSD7A | TUBB3 | ZFP30 |
| PREP | RP11-353N14.1 | SIRT2 | SPTLC3 | TIAF1 | TUBG2 | ZMIZ1 |
| PREX1 | RP11-359M6.1 | SIX1 | SRD5A1 | TIMM8B | TXNRD2 | ZMYND8 |
| PREX2 | RP11-368L12.1 | SLAIN1 | SRGAP2 | TIPARP | UACA | ZNF124 |
| PRH1 | RP11-395N3.2 | SLC16A3 | SRSF1 | TKT | UBA5 | ZNF195 |
| PRICKLE1 | RP11-402L1.11 | SLC16A4 | SRSF3 | TLE3 | UBE2N | ZNF24 |
| PRIM2 | RP11-417J8.3 | SLC16A7 | SSBP3 | TLN2 | UBP1 | ZNF254 |
| PRKAA2 | RP11-435B5.5 | SLC1A2 | ST3GAL3 | TLR1 | UBR7 | ZNF33A |
| PRKCA | RP11-442J21.2 | SLC22A17 | ST3GAL5 | TLR4 | UNC13B | ZNF347 |
| PRKCQ-AS1 | RP11-575F12.3 | SLC22A23 | ST3GAL6 | TMCC1 | UPRT | ZNF385D |
| PRKG1 | RP11-61O11.1 | SLC22A3 | ST6GALC6 | TMEM108 | UQCRB | ZNF430 |
| PRKG2 | RP11-782C8.2 | SLC25A12 | ST8SIA4 | TMEM120A | UQCRC1 | ZNF436 |
| PRLR | RP11-88I18.2 | SLC25A13 | STAC2 | TMEM132A | UQCRH | ZNF444 |
| PROS2P | RP4-665J23.1 | SLC25A23 | STAU2 | TMEM136 | UROS | ZNF454 |
| PRSS12 | RP5-1172A22.1 | SLC25A27 | STK32B | TMEM161B | USP11 | ZNF493 |
| PRTFDC1 | RPS6KA3 | SLC25A29 | STK33 | TMEM173 | USP33 | ZNF506 |
| PRUNE2 | RPS6KA6 | SLC25A5 | STMN1 | TMEM178A | USP46 | ZNF540 |
| PSD3 | RPUSD3 | SLC2A5 | STMN2 | TMEM2 | UTRN | ZNF608 |
| PSMB8-AS1 | RRAGD | SLC35B4 | STON2 | TMEM25 | V1 | ZNF609 |
| PTBP2 | RSRC1 | SLC38A1 | STX12 | TMEM50B | V3 | ZNF618 |
| PTGDS | RUFY3 | SLC38A2 | STX1B | TMEM59L | VAMP4 | ZNF676 |
| PTGFR | RUNX1T1 | SLC38A7 | STX7 | TMEM63B | VAV3 | ZNF680 |
| PTPN13 | S1PR1 | SLC39A10 | STXBP1 | TMEM8A | VCAM1 | ZNF708 |
| PURG | SACS | SLC4A2 | STXBP5 | TMOD2 | VDAC1 | ZNF709 |
| PXYLP1 | SAMD14 | SLC4A4 | SUB1 | TMOD3 | VEZT | ZNF718 |
| PYGB | SARDH | SLC4A7 | SUCLA2 | TMTC2 | VNN1 | ZNF75D |
| RAB13 | SBF2 | SLC6A17 | SULF2 | TMTC4 | VOPP1 | ZNF85 |
| RAB26 | SCG5 | SLC6A8 | SUMO2P18 | TNFRSF19 | VPS13C | ZNRF2 |
| RAB3A | SCML1 | SLC7A6 | SUN1 | TNIK | VPS36 | ZYG11B |
| RAB3C | SCN1A | SLC8A1 | SUSD4 | TNRC6B | VSNL1 | |
| rahiyu | SCN1B | SLC9A9 | SVIL | TNS3 | vubler | |

Table S2. Neuronal genes with PTBP2-mediated splicing events
List of neuronal genes (upregulated in miNs / HAFs; FC ≥ 1.5; ANOVA P < 0.05) exhibiting splicing events associated with PTBP2 during reprogramming (shPTBP2 / shCTL; splicing index ≥ 2; ANOVA P < 0.05).

| Gene name | logFC | Adj.P.Val |
|---|---|---|
| ABCD2 | -1.83E+00 | 9.79E-03 |
| ADAM22 | -1.65E+00 | 9.21E-04 |
| ADAM23 | -1.25E+00 | 1.76E-02 |
| ADCY1 | -2.44E+00 | 5.06E-05 |
| AMOT | -1.20E+00 | 2.04E-02 |
| ANK3 | -1.07E+00 | 7.83E-03 |
| ANKRD6 | -2.24E+00 | 1.15E-05 |
| APBA1 | -1.71E+00 | 3.05E-04 |
| APBB1 | -1.35E+00 | 1.87E-03 |
| APOE | -1.87E+00 | 3.09E-04 |
| ARHGEF7 | -1.03E+00 | 3.36E-03 |
| ASTN1 | -1.64E+00 | 3.52E-04 |
| ATP6V1G2 | -1.33E+00 | 2.66E-02 |
| BAALC | -2.20E+00 | 3.83E-05 |
| BCL7A | -1.24E+00 | 1.66E-03 |
| C14orf132 | -2.08E+00 | 4.63E-05 |
| CALB2 | -3.41E+00 | 7.54E-05 |
| CAMK4 | -1.05E+00 | 2.17E-02 |
| CAMSAP1 | -1.28E+00 | 2.73E-03 |
| CDC7 | -1.98E+00 | 1.97E-03 |
| CERS6 | -2.50E+00 | 6.64E-05 |
| CHRDL1 | -2.53E+00 | 1.88E-05 |
| CKB | -1.40E+00 | 2.44E-03 |
| CLCN4 | -1.32E+00 | 5.14E-03 |
| CNTFR | -1.23E+00 | 2.41E-02 |
| CNTN1 | -1.81E+00 | 9.97E-03 |
| CPE | -1.29E+00 | 2.47E-03 |
| CRISPLD1 | -2.00E+00 | 3.63E-04 |
| CRMP1 | -2.33E+00 | 1.23E-04 |
| CRY1 | -1.30E+00 | 1.28E-03 |
| CSRNP3 | -1.57E+00 | 1.22E-02 |
| CXADR | -1.62E+00 | 2.73E-02 |
| CYFIP2 | -1.14E+00 | 6.06E-03 |
| DCLK1 | -2.28E+00 | 4.58E-05 |
| DCLK2 | -2.34E+00 | 7.21E-06 |
| DPF1 | -2.38E+00 | 8.68E-05 |
| EFR3B | -1.24E+00 | 5.25E-03 |
| ENOX1 | -1.85E+00 | 5.62E-03 |
| FADS2 | -1.73E+00 | 4.46E-04 |
| FAM110B | -1.36E+00 | 7.31E-04 |
| FAM131B | -1.31E+00 | 1.75E-03 |
| FAT3 | -1.51E+00 | 9.85E-04 |
| FGD4 | -1.51E+00 | 8.35E-03 |
| FOXP2 | -2.99E+00 | 3.85E-02 |
| FRY | -1.86E+00 | 5.51E-05 |
| FZD3 | -1.45E+00 | 2.68E-02 |
| GAD1 | -1.28E+00 | 7.76E-04 |
| GDAP1 | -1.34E+00 | 7.20E-03 |
| GPC2 | -1.86E+00 | 1.37E-04 |
| GPD1L | -2.63E+00 | 4.06E-04 |
| GPM6A | -2.35E+00 | 3.96E-04 |
| GPR173 | -1.19E+00 | 1.81E-02 |
| GRIA2 | -1.74E+00 | 2.32E-03 |
| GRIA4 | -1.33E+00 | 6.84E-03 |
| GRM5 | -2.06E+00 | 3.30E-02 |
| GUCY1A2 | -1.54E+00 | 1.75E-02 |
| HMGCS1 | -2.22E+00 | 6.55E-03 |
| IL17RD | -2.54E+00 | 9.18E-06 |
| JPH4 | -1.31E+00 | 6.71E-03 |
| KALRN | -1.45E+00 | 4.23E-04 |
| KCTD12 | -2.29E+00 | 1.44E-04 |
| KIAA0513 | -1.34E+00 | 1.41E-03 |
| KIF5C | -1.34E+00 | 4.06E-03 |
| KLHL25 | -1.32E+00 | 1.78E-03 |
| LRRC4 | -1.37E+00 | 4.46E-03 |
| MAGI3 | -1.48E+00 | 3.88E-04 |
| MAP2 | -1.79E+00 | 5.20E-04 |
| MCM3 | -1.40E+00 | 3.98E-04 |
| METTL7A | -1.80E+00 | 5.74E-05 |
| MGAT4C | -2.81E+00 | 5.42E-03 |
| NCALD | -2.21E+00 | 3.83E-05 |
| NCAM1 | -1.66E+00 | 8.27E-04 |
| NELL2 | -2.45E+00 | 2.15E-05 |
| NFIB | -1.33E+00 | 3.05E-03 |
| OGFRL1 | -2.46E+00 | 2.85E-05 |
| P2RY1 | -2.81E+00 | 1.78E-03 |
| PASK | -1.53E+00 | 4.18E-02 |
| PEG10 | -3.04E+00 | 2.97E-06 |
| PHYHIPL | -2.49E+00 | 9.40E-05 |
| PLCXD2 | -2.58E+00 | 1.68E-04 |
| PODXL2 | -1.18E+00 | 7.68E-03 |
| PPM1L | -1.60E+00 | 2.84E-04 |
| PPP1R12B | -1.76E+00 | 5.18E-05 |
| PRRT2 | -2.31E+00 | 7.95E-05 |
| PTBP2 | -1.70E+00 | 3.41E-02 |
| PTCHD1 | -1.34E+00 | 4.35E-02 |
| RAB3A | -2.17E+00 | 1.87E-04 |
| RALGPS1 | -2.01E+00 | 2.87E-05 |
| RIMS3 | -1.61E+00 | 3.27E-04 |
| RIMS4 | -1.85E+00 | 5.99E-04 |
| RNF144A | -1.18E+00 | 3.01E-03 |
| RNF157 | -1.38E+00 | 1.33E-03 |
| RNFT2 | -1.37E+00 | 4.52E-04 |
| RRAGD | -1.05E+00 | 2.73E-02 |
| RUFY3 | -1.98E+00 | 1.46E-03 |
| SBK1 | -1.75E+00 | 3.09E-03 |
| SCD5 | -1.08E+00 | 3.36E-03 |
| SCN1A | -3.60E+00 | 1.65E-05 |
| SEMA6A | -1.29E+00 | 1.66E-03 |
| SERTAD4 | -2.34E+00 | 1.04E-02 |
| SESN3 | -3.07E+00 | 1.32E-03 |
| SFRP4 | -3.54E+00 | 1.99E-02 |
| SH3BP2 | -1.28E+00 | 1.50E-03 |
| SHF | -2.00E+00 | 4.03E-04 |
| SIAH3 | -2.40E+00 | 4.45E-04 |
| SLAIN1 | -2.35E+00 | 4.30E-05 |
| SLC1A2 | -2.34E+00 | 1.89E-04 |
| SLC4A8 | -1.07E+00 | 3.90E-03 |
| SMAP1 | -1.01E+00 | 1.27E-02 |
| SNAP25 | -1.29E+00 | 2.67E-03 |
| SNCAIP | -3.29E+00 | 9.75E-07 |
| SPARCL1 | -1.35E+00 | 1.21E-02 |
| SPTBN2 | -1.14E+00 | 6.80E-03 |
| SRGAP3 | -1.01E+00 | 1.13E-02 |
| STMN2 | -3.04E+00 | 1.01E-05 |
| STMN3 | -2.03E+00 | 1.84E-04 |
| STRBP | -1.40E+00 | 3.94E-03 |
| SYBU | -2.25E+00 | 5.04E-05 |
| SYT1 | -1.25E+00 | 8.81E-03 |
| THRA | -1.09E+00 | 8.51E-03 |
| THSD7A | -1.32E+00 | 3.91E-03 |
| TMEM108 | -1.26E+00 | 6.35E-03 |
| TMOD2 | -1.76E+00 | 1.57E-03 |
| TMTC2 | -1.29E+00 | 1.11E-03 |
| TNFRSF21 | -1.07E+00 | 2.03E-02 |
| TSPAN14 | -1.11E+00 | 5.54E-03 |
| TSPAN5 | -1.34E+00 | 1.07E-03 |
| TSPAN7 | -2.67E+00 | 8.68E-06 |
| TUB | -1.31E+00 | 8.61E-04 |
| TUBB3 | -2.41E+00 | 1.08E-04 |
| ZNF300 | -1.49E+00 | 6.27E-03 |
| ZNF536 | -1.16E+00 | 1.39E-02 |

Table S3. MiR-124-responsive genes that harbor AGO-enriched peaks in HNs
List of downregulated DEGs (KD / CTL; $Log_2FC \geq 1$; adj. P.value > 0.05) in HNs with miR-124 and ELAVL3 knockdown that harbor AGO-enriched peaks determined by HITS-CLIP in miNs

| | |
|---|---|
| ABCD2 | PHYHIPL |
| ADAM22 | PLCXD2 |
| ADAM23 | PPM1L |
| ADCY1 | PPP1R12B |
| ANK3 | PTBP2 |
| APBA1 | RALGPS1 |
| ARHGEF7 | RIMS3 |
| ASTN1 | RNF144A |
| BAALC | RUFY3 |
| BCL7A | SCD5 |
| CAMSAP1 | SCN1A |
| CLCN4 | SEMA6A |
| CNTN1 | SESN3 |
| CPE | SLAIN1 |
| CXADR | SLC1A2 |
| CYFIP2 | SLC4A8 |
| DCLK1 | SMAP1 |
| DCLK2 | SNAP25 |
| EFR3B | SPARCL1 |
| ENOX1 | SRGAP3 |
| FAM110B | STRBP |
| FAM131B | SYT1 |
| FAT3 | TMEM108 |
| FGD4 | TMTC2 |
| FRY | TNFRSF21 |
| FZD3 | TSPAN5 |
| GDAP1 | TSPAN7 |
| GPD1L | TUB |
| GPM6A | ZNF536 |
| GRIA2 | |
| GRIA4 | |
| GRM5 | |
| GUCY1A2 | |
| HMGCS1 | |
| JPH4 | |
| KALRN | |
| KCTD12 | |
| KIAA0513 | |
| KIF5C | |
| MAGI3 | |
| MAP2 | |
| METTL7A | |
| MGAT4C | |
| NCALD | |
| NCAM1 | |
| NELL2 | |
| NFIB | |
| PEG10 | |

Table S4. MiR-124-responsive genes predicted to be bound by nELAVLs in HNs
List of miR-124-responsive genes predicted to be bound by nELAVLs (Scheckel et al., 2016) in HNs.
